# Characterization of sleep in a mouse model of CLN3 disease revealed sex-specific sleep disturbances

**DOI:** 10.1101/2024.05.24.595712

**Authors:** Kelby M. Kane, Diane Iradukunda, Christopher J. McLouth, Landys Z. Guo, Jun Wang, Anjana Subramoniam, Dillon Huffman, Kevin Donohue, Bruce F. O’Hara, Sridhar Sunderam, Qing Jun Wang

## Abstract

The neuronal ceroid lipofuscinoses (NCLs) are a group of recessively inherited neurodegenerative diseases characterized by lysosomal storage of fluorescent materials. CLN3 disease, or juvenile Batten disease, is the most common NCL that is caused by mutations in the *Ceroid Lipofuscinosis, Neuronal 3* (*CLN3*) gene. Sleep disturbances are among the most common symptoms associated with CLN3 disease, yet this is understudied and has not been delineated in an animal model of the disease. The current study utilized a non-invasive, automated piezoelectric motion sensing system (PiezoSleep) to classify sleep and wakefulness in a *Cln3^ϕ..ex^*^1–6^*^/ϕ..ex^*^1–6^ (*Cln3*KO) mouse model and age- and sex-matched wild-type (WT) controls. The sleep-wake classification by PiezoSleep was found to be about 90% accurate when validated against simultaneous gold standard polysomnographic recordings including electroencephalography (EEG) and electromyography (EMG) in a small cohort of WT and *Cln3*KO mice. Our large cohort PiezoSleep study reveals sleep abnormalities during the light period (LP) in male *Cln3*KO mice compared to WT male, and more subtle differences in *Cln3*KO female mice throughout the dark period (DP) compared to WT female, recapitulating sleep abnormalities seen in CLN3 disease patients. Our characterization of sleep in a mouse model of CLN3 disease contributes to a better understanding of the sleep disturbances commonly reported for CLN3 disease and other NCLs, which will facilitate the development of new disease treatment and management strategies.

## Introduction

### CLN3 disease

CLN3 disease, also called juvenile Batten disease or juvenile neuronal ceroid lipofuscinosis (JNCL), is among the group of rare lysosomal storage disorders that share the characteristics of intracellular accumulation of lipofuscin and collectively are the most common inherited neurodegenerative disorders of childhood. CLN3 disease is caused by mutations in the *CLN3* gene, with the most common mutation being a 966-bp deletion spanning exons 7 and 8 on human chromosome 16p12 ^1^. The *CLN3* gene encodes a 438 amino acid transmembrane protein that is primarily localized in endosomal/lysosomal compartments and is expressed in multiple human tissues ^2^. Studies using animal and cell model systems of CLN3 disease have shed light on a wide range of roles that CLN3 protein may participate in, including: intracellular trafficking ^3–6^, autophagy ^7–9^, Ca^2+^ levels ^10^, lysosomal egress ^11^, and lysosomal pH ^12^. However, the exact pathway(s) that CLN3 is involved in are still debatable, which may contribute to the lack of treatments for this disease despite of current development of potential interventions such as gene therapy ^13,14^. Clinically, the most common (in 80% of patients) and often an early symptom is functional vision loss, the onset of which occurs around 5-6 years of age ^15^. Signs of cognitive impairment and behavior changes typically appear approximately two years later. In teenage years, loss of memory and independent skills such as mobility, feeding, and communication are present. Patients are bedridden by their 20s, and death typically occurs in the third decade of life ^16^.

### Sleep and CLN3 disease

Besides the clinical symptoms mentioned above, the majority of CLN3 disease patients experience poor sleep ^17^. Sleep is a basic physiological necessity essential for healthy brain function, but people can fall short of their sleep requirement due to environmental factors, lifestyle choices, and changes to the body, such as pain. Problems with sleep reported in CLN3 disease patients include difficulty falling asleep, nocturnal awakenings, nightmares, and restless sleep ^18^. In the typical time course relative to the onset of CLN3 disease, sleep disturbances start to occur at an average age of 8-10 years as a clinical marker ^16^. Heikkilä et al. found that average sleep percentage during the night was lower in patients with CLN3 disease compared to control subjects, and higher in the daytime ^19^. Interestingly, the majority of the patients had normal melatonin, cortisol, and body temperature rhythms, indicating that the abnormal sleep-wake cycles could not be explained by failure of the circadian regulatory system ^19^. A more recent 2016 study in a cohort of 28 CLN3 disease patients showed that 96% of patients experienced sleep disturbances ^17^. When considering the heterogeneous groups of NCL including CLN1, CLN2, CLN3, and CLN5 diseases as a whole, the age of onset of sleep disturbances is moderately associated with the onset of both vision loss and seizures, although each individual form of NCL showed no significant correlations between these factors ^17^.

Sleep can be divided into rapid-eye movement (REM) sleep and non-REM (NREM) sleep. NREM sleep in humans can be further parsed by the frequency and amplitude of oscillations in the EEG into three sub-stages: NREM stage 1 (N1), stage 2 (N2), and stage 3 (N3). N1 is light sleep with EEG waves highest in frequency and lowest in amplitude. N2 is a deeper stage of sleep with the EEG baseline interrupted by sleep spindles and K complexes. N3, also called slow wave sleep, is the deepest stage of sleep with the EEG dominated by large amplitude slow oscillations (i.e., delta waves) ^20^. Kirveskari et al. compared sleep structure in 28 CLN3 disease patients with healthy control subjects using polysomnography ^21^. They reported that total sleep time, sleep efficiency index, and percentages of REM and N2 sleep were significantly lower in the patients, while percentages of N1 and N3, and the number of nocturnal awakenings were significantly increased ^21^, suggesting sleep disturbances.

### Brief overview of the work

The past sleep studies and case reports only provide a limited window into the sleep disturbances that CLN3 patients experience. In particular, studies utilizing questionnaires may be subject to bias and inaccuracy as they are completed by family members, and the broad age range and age of disease onset in patient cohorts are elements that add variability into the data and are difficult to control. Case reports aside, no sleep study has yet investigated the mechanism behind disordered sleep in CLN3 disease. This is where preclinical models can play a critical role. Here we report the first study utilizing CLN3 disease mouse models to provide insight into sleep disturbances associated with the disease. Specifically, we utilized *Cln3^ϕ..ex^*^1–6^*^/ϕ..ex^*^1–6^ mice with homozygous deletion of *Cln3* exons 1-6 (abbreviated as *Cln3*KO hereafter) ^22^. Utilizing a piezoelectric motion sensing system ^23,24^, we continuously monitored sleep over several days and analyzed percentage sleep, mean sleep bout length, and mean top 10% sleep bout length. To better understand when these sleep differences occur, we also analyzed the 24-hour time course of sleep patterns. Our results show sex-dependent patterns of sleep disturbances in *Cln3*KO mice. Of note, we also obtained simultaneous EEG/EMG recordings alongside piezoelectric measurements from a separate and small cohort of female *Cln3*KO and WT mice to analyze sleep composition and validate the accuracy of sleep scoring by the piezoelectric system.

## Methods

### Animal Model

Both *Cln3*KO (B6.129S6-*Cln3^tm1Nbm^*/J, #029471; *Cln3^τ1.ex^*^1–6^*^/τ1.ex^*^1–6^; originally made by Mitchison *et al* ^22^) and sex- and age-matched C57BL/6J (#000664) WT controls were purchased from The Jackson Laboratory. This *Cln3KO* strain was backcrossed to C57BL/6J for 10 generations by David Pearce before donating to Jax. Genetic monitoring was carried out by Transnetyx based on miniMuGA array ^25^ on a subset of the *Cln3KO* animals and confirmed that these animals are congenic on C57BL/6J background except for the region at chromosome 7 where mouse *Cln3* gene resides prior to deleting exons 1-6.

### Study Design

All animal procedures were performed under approved IACUC protocols (#2016-2313 and #2017-2838) at the University of Kentucky. Before the study, mice were housed in a 14h:10h light-dark (LD) cycle environment. A noninvasive sleep monitoring system using piezoelectric “piezo” motion sensors (PiezoSleep, Signal Solutions LLC, Lexington, KY; www.sigsoln.com) was validated in a small number of *Cln3*KO (n=3) and WT (n=4) female mice aged 8-14 months against gold standard invasive EEG/EMG recordings and then used to score baseline sleep using automated PiezoSleep software in a large cohort of male and female *Cln3*KO and WT mice of various ages (2, 4, 6, and 8-10 months old; **Table 1**).

**Table 1.** Numbers of mice used at different ages in the large cohort PiezoSleep studies. Of note, for some of the mice, PiezoSleep was assessed at multiple ages.

| Ages | 2 months | 4 months | 6 months | 8-10 months |
| --- | --- | --- | --- | --- |
| Male WT | 9 | 9 | 9 | 9 |
| Male *Cln3*KO | 9 | 7 | 7 | 6 |
| Female WT | 11 | 4 | 6 | 5 |
| Female *Cln3*KO | 11 | 4 | 5 | 6 |

### Surgical procedures for EEG/EMG

Mice were surgically instrumented for EEG/EMG recording using procedures previously described ^23,26^. Briefly, animals were given oral analgesia (Carprofen, 10 mg/kg s.c.) and shaved to expose the scalp under 2.5% isoflurane anesthesia. They were then secured in a recumbent position using ear bars on a stereotactic frame. A scalp incision was made along the midline from behind the eyes to the neck and the exposed skull swabbed with povidone-iodine solution and 70% isopropyl alcohol. A prefabricated rectangular EEG/EMG head mount (8201-SS, Pinnacle Tech., Lawrence, KS) was glued to the skull centered between bregma and lambda and secured at the four corners with stainless steel bone screws (8209 and 8212, Pinnacle Tech.) painted with silver epoxy to enhance conductivity. The screws served as frontal and parietal EEG electrodes with common reference and ground. Bipolar stainless-steel wires at the rear of the head mount were inserted into the nuchal muscle to record muscle tone (EMG), which differentiates sleep and wakefulness. The incision was closed with 2-3 sutures, and the head mount sealed with dental cement to prevent infection. The animals were given analgesia for the next three days. The implant was allowed to set for two weeks before suture removal.

### Simultaneous EEG/EMG and PiezoSleep data acquisition

For sleep recording, the animals were moved to custom-built cages with open ceilings and free access to food and water on a 12h:12h LD cycle. EEG and EMG signals were recorded by a data acquisition system (DACS No. 8206, Pinnacle Tech.) through a preamplifier (10x or 100x, No. 8202, Pinnacle Tech.) plugged into the head mount and suspended from above by a slipring commutator arrangement. Signals were sent through a lowpass antialiasing filter (25 Hz cutoff for EEG and 100 Hz for EMG) by the DACS, sampled at 400 Hz, and saved on a computer hard drive using Pinnacle’s Sirenia 1.7.2 acquisition software. Additionally, a piezoelectric motion film sensor (Signal Solutions) placed on the cage floor under a plastic liner recorded piezoelectric signals synchronized with the EEG and EMG as an auxiliary channel on Sirenia.

### Sleep classification

Of several multi-day recordings, those with a minimum length of five continuous days were selected. Ignoring the first two days to allow for acclimatization to the new cage environment, recordings from days 3-5 were used for analysis. In each 24-hour recording, sleep was scored separately from the EEG/EMG and piezo signals in non-overlapping 4-second epochs (**Figure 1**) and compared with each other. The piezo signal separated wakefulness from sleep based on motion, the EMG separates wakefulness from sleep based on muscle tone, and the EEG further distinguishes NREM and REM stages within sleep based on cortical neural activity. NREM sleep is characterized by high delta power (0.5-4 Hz) and REM sleep by high theta power (6-9 Hz). Signal epochs were first classified as sleep or wake based on mean-squared EMG power using a two-component Gaussian mixture model (GMM). Then the sleep epochs were further classified as NREM or REM sleep using another GMM fitted to the EEG delta/theta EEG power ratio and high/low frequency band power ratio (i.e., ratio of 9-80 Hz to 0.5-9 Hz power). The cluster with the higher mean delta/theta power ratio was labelled as NREM and the other as REM (**Figure 1**). Furthermore, a sleep-wake decision statistic was extracted from the piezo signal using SleepStats software (Signal Solutions) and a two-component GMM also fitted to it to classify signal epochs as sleep or wake for further analysis.

**Figure 1.**
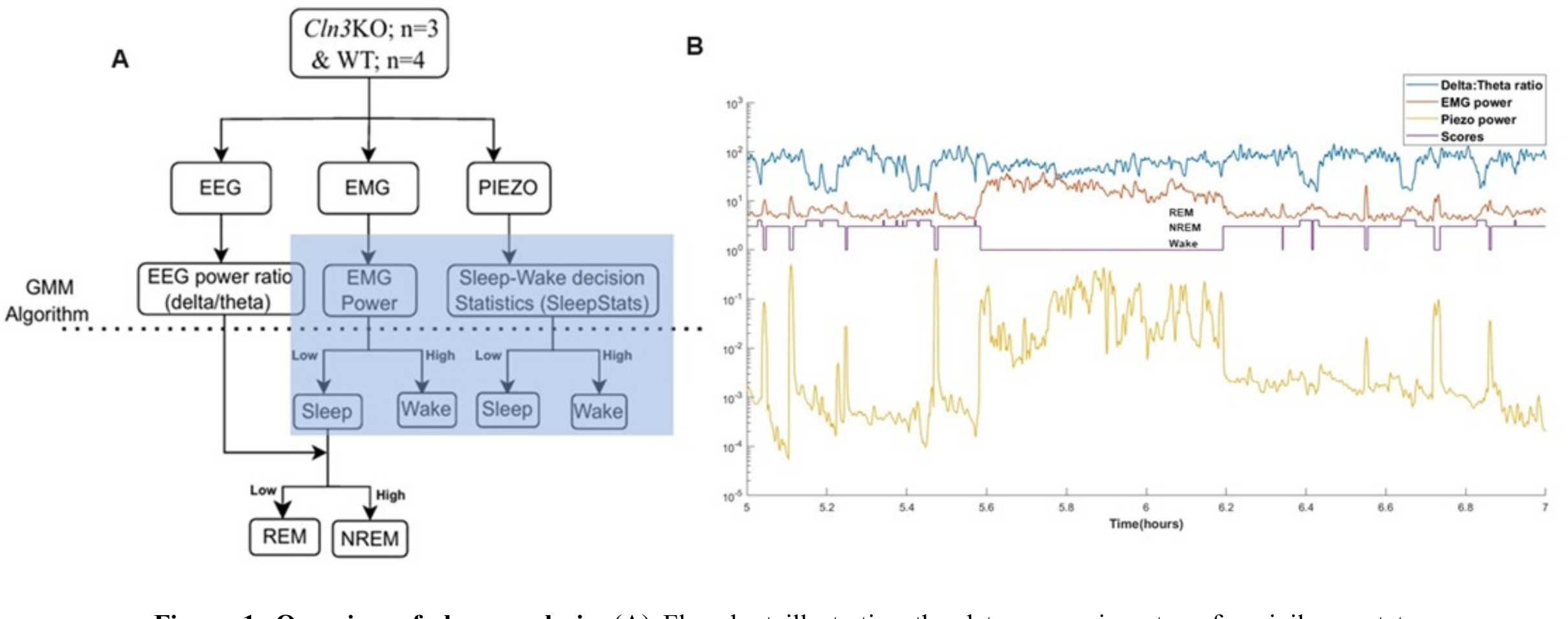
Overview of sleep analysis. (**A**) Flowchart illustrating the data processing steps for vigilance state assessment; (**B**) Snapshot of signal features (EEG delta to theta power ratio, EMG power, piezo power) and corresponding sleep scores (Wake, NREM and REM sleep).

### Statistical analysis for simultaneous EEG/EMG and PiezoSleep

The noninvasive piezo sleep scores were validated against the EEG/EMG scores as a reference to benchmark their accuracy. The correlation between the PiezoSleep decision statistic and mean-squared EMG power was tested using the Spearman rank coefficient. Piezo-based scoring accuracy was quantified against EMG-based scores using conventional detection metrics: namely, sensitivity, specificity, and precision. Sleep-wake scores were used to estimate typical metrics (percent time, mean bout length) in 30-minute intervals over the 24-hour cycle and separately for the light (ZT 0-12h) and dark (ZT 12-24h) periods of the day. Piezo-based sleep metrics were compared against those derived from the EMG using the Mann-Whitney U test in GraphPad Prism V10. Data are presented as mean ± SEM, and a type I error probability of 0.05 was considered statistically significant. Finally, percent time and mean bout length for NREM, REM, and total sleep were compared for *Cln3*KO and WT mice using the Mann-Whitney U test in GraphPad Prism V10 (**Table 2**).

**Table 2.** Tests of Significance (Mann-Whitney) for the Sleep Stage Differences Between *Cln3*KO and WT Mice during the Light Period (LP) and Dark Period (DP).

|  | Sleep% (Figure 4)<br>p-value | Mean Bout Length (Figure 5)<br>p-value |
| --- | --- | --- |
| LP | 0.8571 | 0.8571 |
| DP | 0.4 | >0.9999 |
| NREM LP | >0.9999 | 0.6286 |
| NREM DP | 0.4 | 0.8571 |
| REM LP | 0.6286 | 0.4 |
| REM DP | 0.6286 | 0.0571 |

### Large cohort PiezoSleep data acquisition and processing

The mice listed in **Table 1** were singly housed in the PiezoSleep Adapt-a-Base cage system (Signal Solutions, Lexington, KY) in a room with a 12h:12h LD cycle; food and water were available *ad libitum*. The motion sensor system includes piezo disc sensors, which, like in the piezo film sensor system described previously ^27,28^, are placed at the bottom of the cage to generate a “PiezoSleep” signal that helps distinguish between sleep and wakefulness on the basis of subtle animal movements. The system automatically scores sleep and wakefulness, where sleep states are characterized by breathing-related quasi-periodic signals with low variation in amplitude, and active or resting states during wakefulness are detected as irregular transients and high amplitude pressure variations from the mouse moving and shifting its weight ^28^.

Signals collected through the PiezoSleep system were processed by the SleepStats software package (Signal Solutions; v2.181). Unlike humans who most commonly exhibit monophasic sleep with sleep duration between 7 to 8 hours (in 80-120 minute NREM-REM cycles), rodent sleep is fragmented and polyphasic throughout the light phase and dark phase, with more sleep and slightly longer bouts in the light phase^29^. Therefore, with the minimum sleep bout length set as 4 s, sleep bouts were identified and accumulated over 30-minute bins to yield three sleep parameters: 1) percent time in sleep; 2) mean sleep bout length; and 3) mean top 10% longest sleep bout length.

### Large cohort PiezoSleep data analyses and statistics

As it took animals 1-2 days to acclimate to the PiezoSleep cages and LD cycle of the test room, only data from days 3 to 5 (full days starting from ZT0 to ZT24) were used in the statistical analysis. Two types of statistical analyses were performed, one for 30-minute time intervals and the other for 12-hour averaged light- and dark-period sleep measurements. General linear mixed models were used to analyze sleep data to properly account for repeated measures. Specifically, the study design utilized two repeated measures: mouse age (categorized as 2, 4, 6, and 8-10 months) and time of day (48 consecutive 30-minute intervals starting at ZT0). The primary outcomes were sleep percent, mean sleep bout length, and mean top 10% sleep bout length, averaged over every 30-minute interval and then over study days 3 through 5. These outcomes were predicted by the two within-subject factors (age and time of day) and two between-subjects factors: genotype (WT *vs. Cln3*KO) and sex (male *vs.* female). Models were initially estimated with all main effects and higher-order interactions, and non-significant four-way interactions were removed in favor of a more parsimonious model. Significant three-way interactions were followed up with tests of simple effects. When exploring significant interactions, the Benjamini-Hochberg procedure was used to control the false discovery rate (FDR) at 0.05 within each outcome^30^. The 12-hour averaged light- and dark-period analyses also used a general linear mixed model, but time of day was no longer a within-subject factor since these measures were averaged into two separate outcomes. We additionally included early dark phase (ZT12 – ZT18) as an outcome. All data were analyzed using PROC MIXED from SAS version 9.4 (SAS Institute Inc, Cary, NC).

## Results

### Validation of the PiezoSleep system

#### Feature correlation and sleep scoring accuracy

The PiezoSleep motion sensor-based system scores sleep with over 90% accuracy as verified by human observation and simultaneous EEG recordings in different strains of mice ^23,24,31^. We first sought to validate the PiezoSleep system, which is a more recent variation of the technology, to make sure it would be accurate for the specific *Cln3*KO and WT mice to be used in the large cohort study. We found strong and consistent correlations across both WT and *Cln3*KO mice (combined n=7) between the piezo and EMG features that were used as the bases for discriminating between sleep and wake states (**Figure 2**). The Spearman rank correlation coefficient between the PiezoSleep decision statistic and mean-squared EMG power, both of which were used to distinguish between wakefulness and sleep states, averaged about 72% during the light period (LP) and 70% during the dark period (DP) (**Figure 2A**), indicating strong concordance between the two measurement methods regardless of genotype and time of the day. A representative scatter plot (**Figure 2B**) further demonstrates the relationship between these EMG and piezo features. This analysis provides additional evidence of agreement between Piezo and EMG measurements in discriminating sleep and wake, reinforcing the reliability of both methods. The potential accuracy of the piezo statistic when compared against a threshold to differentiate sleep and wake states was assessed against the EMG-based scores using a receiver operating characteristic (ROC) analysis. The ROC curve charts the effect of a chosen threshold on the sensitivity and specificity of sleep detection with respect to the EMG scores. The area under the ROC curve (AUC) reflects the potential accuracy of the detection feature, with values close to 1 being more desirable. As evidenced by a representative ROC curve (**Figure 2C**), our data show consistent and high mean AUC values of 0.940 for DP and 0.937 for LP, indicating accurate discrimination between sleep and wake states. Together, these results confirm the reliability of the PiezoSleep system in assessing sleep architecture across genotypes and LD cycle with 90% or higher sensitivity and specificity at the optimal threshold corresponding to the knee (top left corner) of the ROC curve.

**Figure 2.**
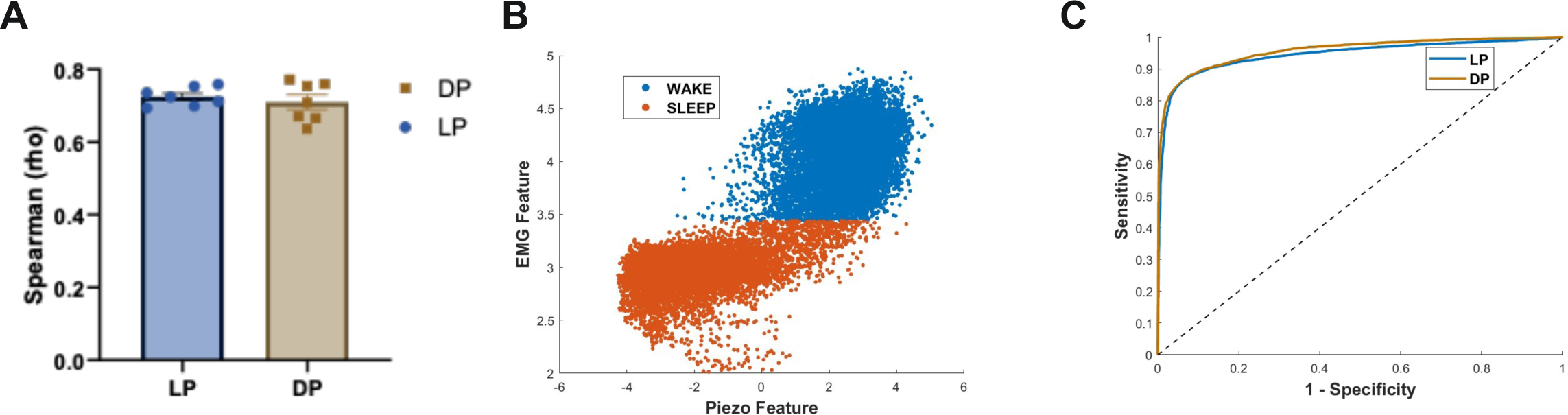
Validation of noninvasive sleep analysis method. (**A**) Spearman rank correlation coefficient shows strong agreement between the PiezoSleep decision statistic and mean-squared EMG power during light period (LP; average rho=0.72) and dark period (DP; average rho=0.71), based on data from the 7 mice in the verification cohort. *Cln3*KO mice are outlined in black; (**B**) A representative scatter plot of EMG mean squared power and piezo sleep-wake decision statistic features, showing differentiation of wakefulness and sleep, from a 14-month-old-female *Cln3*KO mouse; (**C**) A representative receiver operating characteristic (ROC) analysis with high AUC values (0.937 for LP and 0.940 for DP) demonstrates accurate discrimination between sleep and wake states using PiezoSleep decision statistic with EMG as a reference across the 24-hour cycle (LP and DP) from a14-month-old-female *Cln3*KO mouse.

#### Comparison of Piezo and EEG sleep metrics

The percent time spent in sleep (Sleep%) and the mean bout length (MBL) for each mouse in the validation cohort (n=7; 4 WT and 3 *Cln3*KO) were calculated from the piezo and EMG recordings and compared. Our results show consistency in Sleep% calculated from the piezo and EMG recordings during both LP and DP, with an absolute error less than about 2 % for 5 of 7 animals (**Figure 3A**). In contrast, sleep MBL calculated from EMG recordings were consistently longer compared to values calculated from piezo recordings during both LP (p=0.016) and DP (p=0.006) (**Figure 3B**). A closer examination revealed that the piezo signal was more sensitive to brief arousals in sleep than the EMG. This may be attributed to differences in the location of measurement, with the EMG being closer to the head and therefore sensitive to head and neck movements while the piezo sensor is in contact with the animal’s ventral surface and is therefore more sensitive to changes in ventilation or twitches in the lower body as may occur during REM sleep. In effect, the piezo adds more brief arousals to the sleep record and therefore breaks up sleep into shorter bouts. If we progressively remove brief arousals starting with the shortest ones, the piezo and EMG estimates of MBL converge after arousals under 30s are dissolved (**Figure S1**). This demonstrates that the two measurements are consistent with each other.

**Figure 3.**
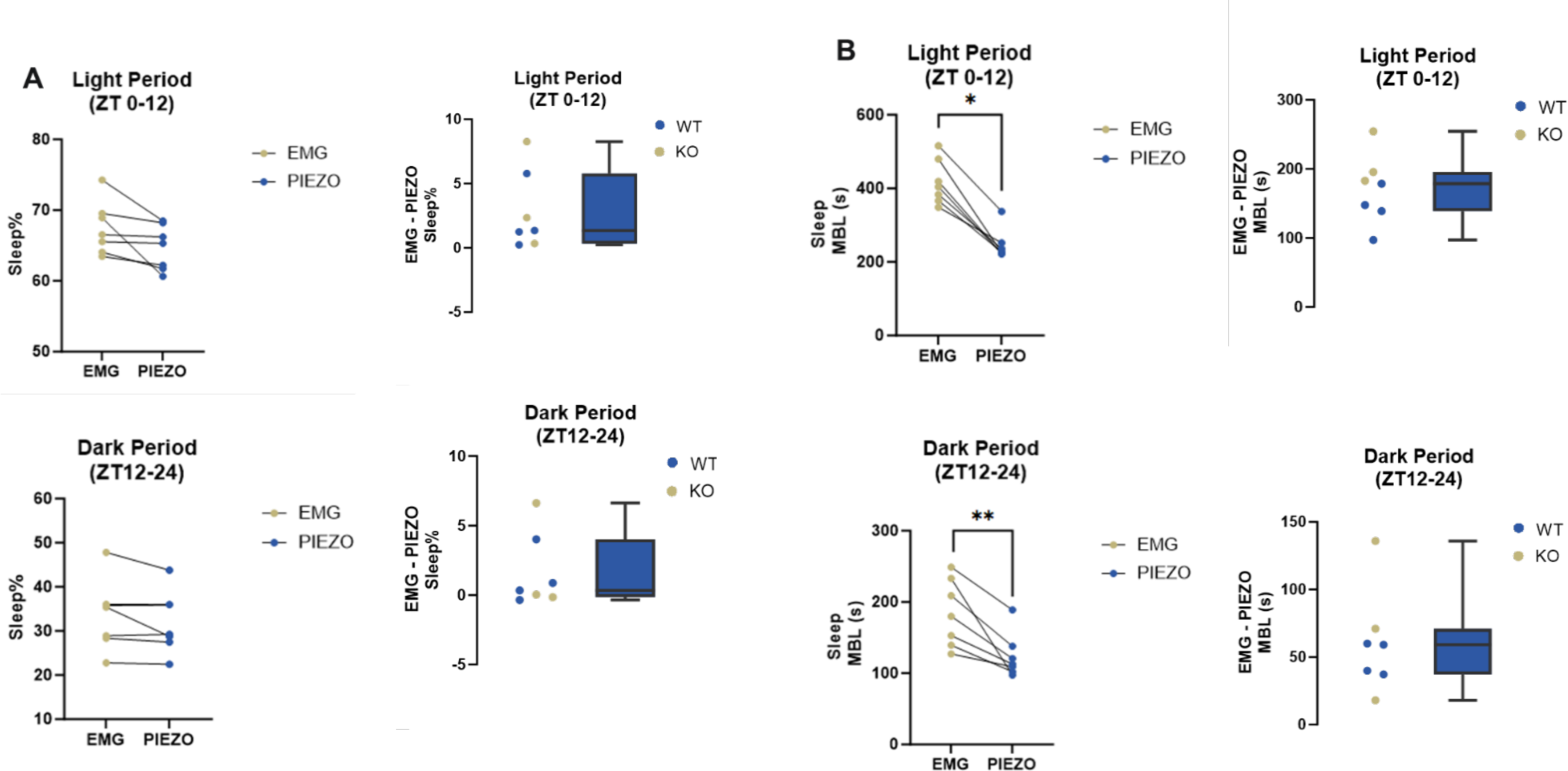
Comparison of sleep metrics between piezo and EEG recordings in female mice (n=7; 3 *Cln3*KO and 4 WT). (**A**) shows the consistency in percent time and associated differences (EMG minus piezo) obtained from both piezo and EMG recordings during LP and DP; (**B**) represents sleep MBL during both LP and DP, with differences highlighted by the absolute error plots. *p<0.05, **p<0.001, ****p<0.0001.

#### Comparison of EEG/EMG-based sleep metrics for Cln3KO and WT mice

We conducted EEG/EMG analyses to score all three states of vigilance (wake, NREM, REM) for WT (n=4) and *Cln3*KO (n=3) mice. While this sample size is too small to reach statistically significant conclusions (**Table 2**) for either Sleep% (**Figure 4**) or MBL (**Figure 5**), MBL for REM sleep in DP in *Cln3*KO tended to be longer than that in WT (**Figure 5F**; p=0.0571, **Table 2**). These differences were borne out by the 24-hour profiles in these sleep metrics (data not shown). Results from this validation cohort study suggest that analysis of genotype differences in sleep is warranted in a cohort with larger sample size.

**Figure 4.**
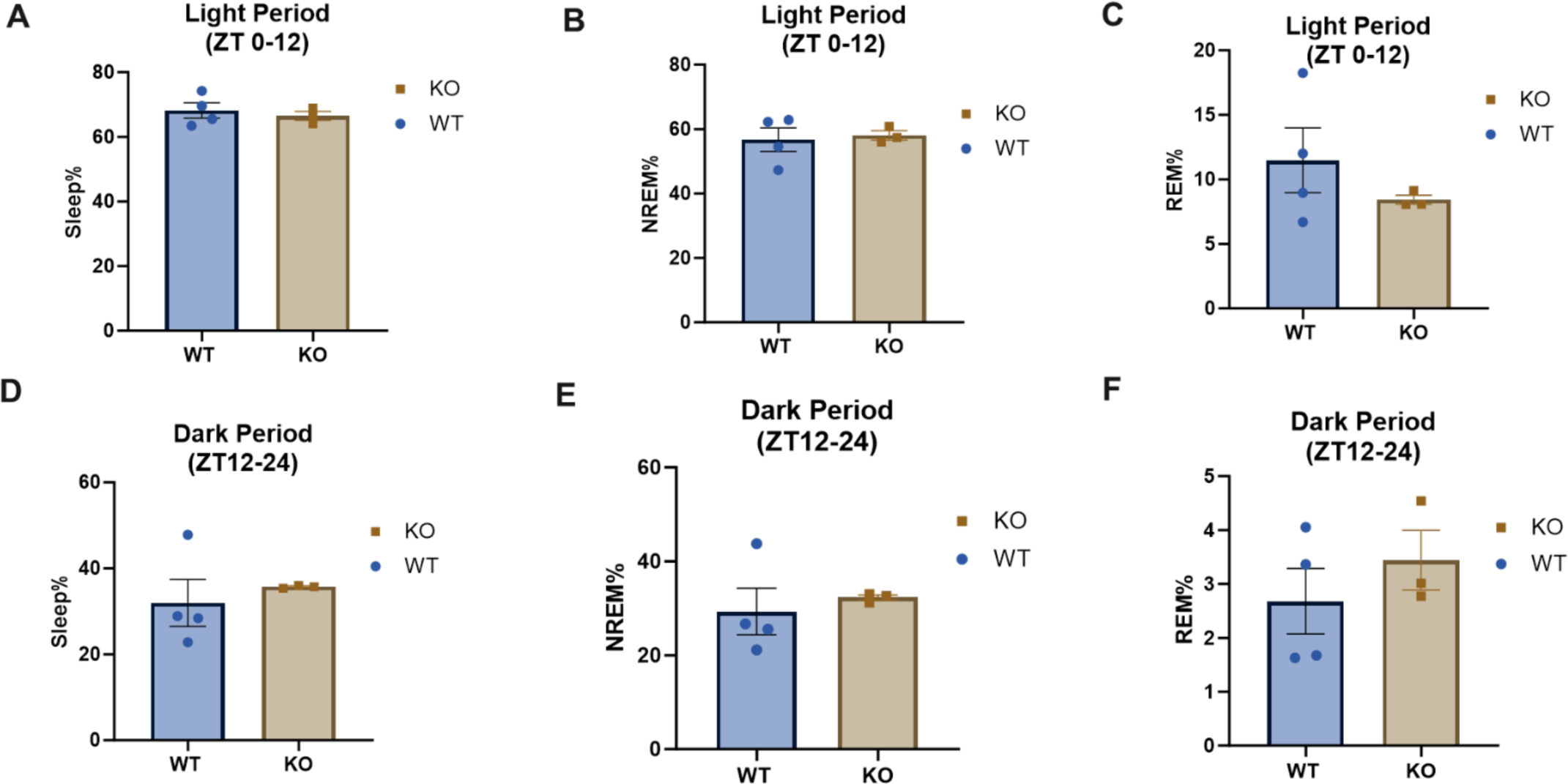
Comparison of EEG/EMG-based sleep stage metrics of Sleep% between *Cln3*KO and WT mice during LP and DP (n=7; 3 *Cln3*KO and 4 WT). **(A)** and **(D)** represent % sleep during LP and DP; (**B**) and (**E**) represent %NREM sleep during the LP and DP; (**C**) and (**F**) represent REM% sleep during the LP and DP. Mann-Whitney U tests were performed, and p-values recorded in **Table 2**.

**Figure 5.**
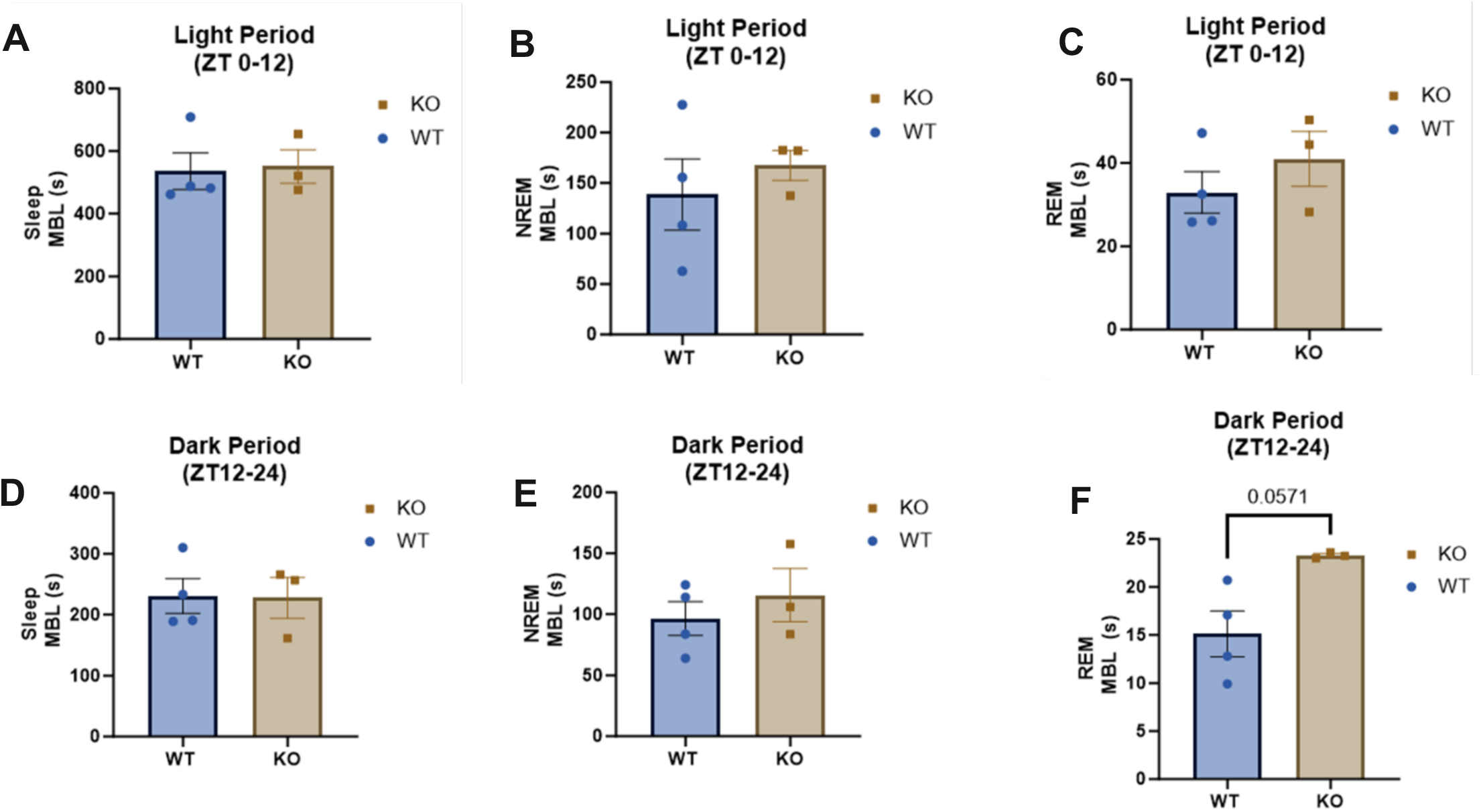
Comparison of EEG/EMG-based sleep stage metrics of Sleep MBL in *Cln3*KO and WT mice during LP and DP. Mann-Whitney tests were performed. Although not statistically significant, MBL is consistently slightly greater for *Cln3*KO (n=3) than WT mice (n=4), particularly for REM MBL (Figure 5F) during DP (p=0.0571; **Table 2**).

### PiezoSleep Analysis of a large cohort of WT and Cln3KO mice

#### Percent sleep abnormalities in Cln3KO mice

We first looked at 24-hour time course of percent sleep of WT and *Cln3*KO male and female mice at all ages. There were no effects involving genotype that were significant at any time points across the 24-hour period (**Table 3**, **Figure 6**). However, there appeared to be trends that male *Cln3*KO had lower sleep percent compared to male WT in the light period (LP, i.e., ZT 0-12; **Figure 6B**) and female *Cln3*KO had higher sleep percent compared to female WT in the dark period (DP, i.e., ZT 12-24; **Figure 6C**). Therefore, we analyzed light and dark periods separately. Because there seemed to be more pronounced differences in the earlier hours of the dark period (**Figure 6A**), we decided to also analyze the first 6 hours of the dark period, herein termed the early dark period (EDP, i.e., ZT 12-18). Our analyses show that genotype by sex interaction for percent sleep during the LP was significant (p = 0.0063; **Table 4**). This was followed up with tests of simple effects, which revealed that male (but not female) *Cln3*KO indeed had lower sleep percent compared to male WT (**Figure 7A**, p = 0.0024). Interestingly, during the EDP, despite non-significant genotype by sex interaction, there was a trend of increased percent sleep in *Cln3*KO with mixed sex (p = 0.0581, **Table 4**; **Figure 7C**).

**Table 3.** Overall Tests of Significance for the Sleep Time Course Data

| *Effect* | **Sleep Percent** |  | **Mean Sleep Bout Length** |  | **Mean Top 10% Sleep Bout Length** |  |
| --- | --- | --- | --- | --- | --- | --- |
|  | **F-value** | **p-value** | **F-value** | **p-value** | **F-value** | **p-value** |
| *Time of Day (T)* | 78.66 | <.0001 | 64.26 | <.0001 | 99.42 | <.0001 |
| *Mouse Age (A)* | 15.38 | <.0001 | 12.97 | <.0001 | 7.33 | 0.0003 |
| *Genotype (G)* | 0 | 0.9844 | 0.72 | 0.4011 | 1.11 | 0.299 |
| *Mouse Sex (S)* | 22.88 | <.0001 | 16.78 | 0.0001 | 11.18 | 0.0017 |
| *T by A* | 1.98 | <.0001 | 1.52 | <.0001 | 1.82 | <.0001 |
| ***T by G*** | 1.22 | 0.1474 | 1.8 | **0.0007** | 2.33 | <.0001 |
| *T by S* | 5.05 | <.0001 | 2.48 | <.0001 | 3.07 | <.0001 |
| *A by G* | 0.76 | 0.517 | 1.15 | 0.3289 | 0.45 | 0.7187 |
| *A by S* | 1.33 | 0.2644 | 1.89 | 0.1296 | 1.14 | 0.3408 |
| ***G by S*** | 2.64 | 0.1109 | 5.29 | **0.0253** | 4.59 | 0.038 |
| *T by A by G* | 0.9 | 0.7917 | 1.1 | 0.1952 | 1.03 | 0.3853 |
| ***T by A by S*** | 1.45 | **0.0005** | 1.24 | **0.0286** | 1.63 | **<0.0001** |
| ***T by G by S*** | 0.69 | 0.9468 | 1.01 | 0.4603 | 1.48 | **0.0182** |
| *A by S by G* | 0.36 | 0.7816 | 1.62 | 0.1825 | 0.74 | 0.5344 |

Red bold: Significant genotype dependent comparisons
Blue bold: Significant non-genotype dependent comparisons

**Figure 6.**
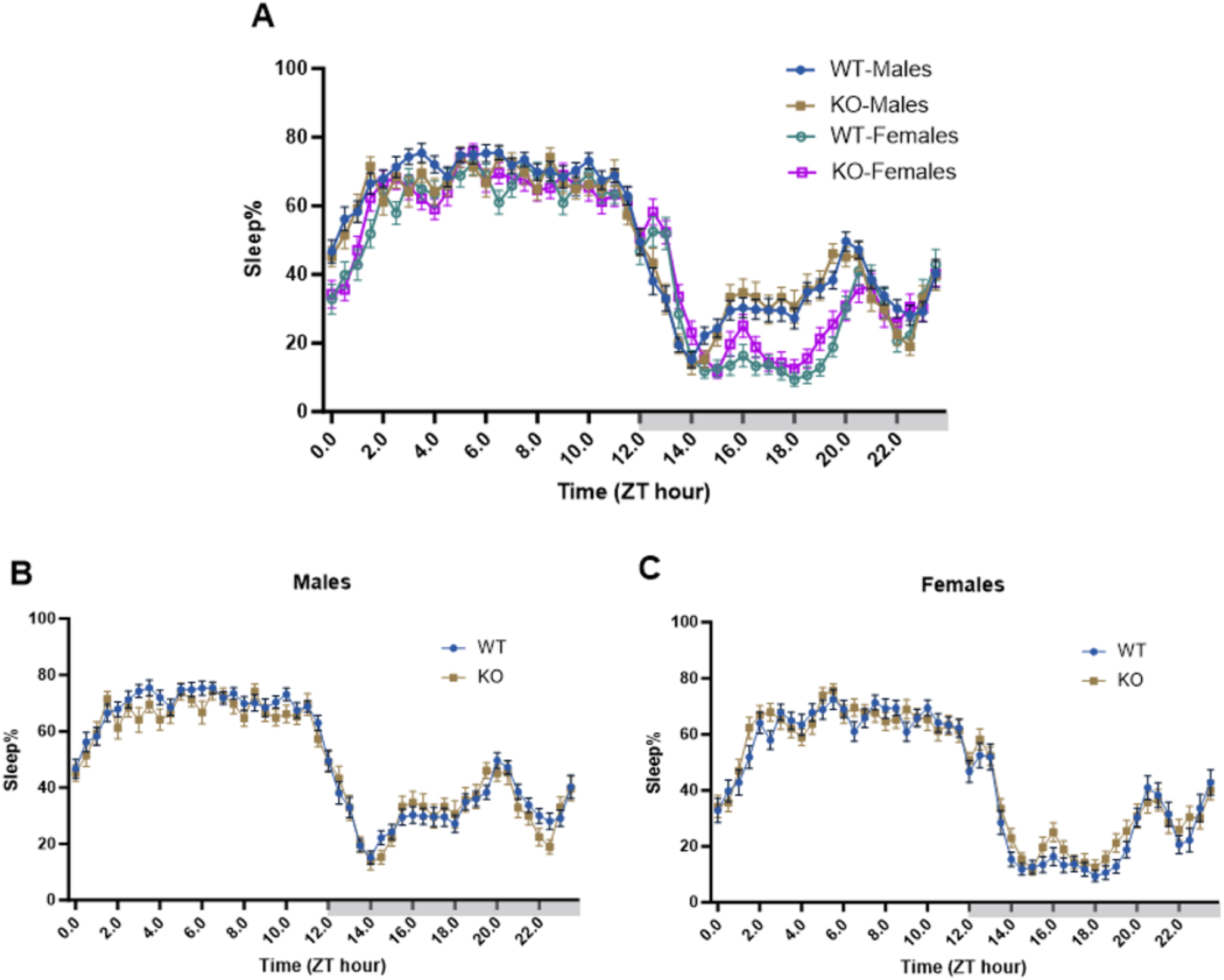
24-hour time course of WT and *Cln3*KO male and female mice. Graphs depict 24-hour sleep traces with sleep percentages every 30 minutes, averaged from days 3-5. Groups included several different ages at the onset of Piezo Sleep acquisition (2 months N= 9 for each genotype; 4 months = 6 months = 9 WT, 7 *Cln3*KO; 8-10 months = 9 WT, 6 *Cln3*KO) and females of several different ages (2 months N=11 for each genotype; 4 months N= 4 WT, 4 *Cln3*KO; 6 months = 6 WT, 5 *Cln3*KO; 8-10 months = 5 WT, 6 *Cln3*KO). (**A**) represents male and female WT and *Cln3*KO mice plotted together on the same graph; (**B**) and (**C**) depict WT and *Cln3*KO mice separated by sex. Each data point is represented as mean ± SEM (Male WT n= 31; male *Cln3*KO n= 22; female WT n =26; female *Cln3*KO n= 26).

**Figure 7.**
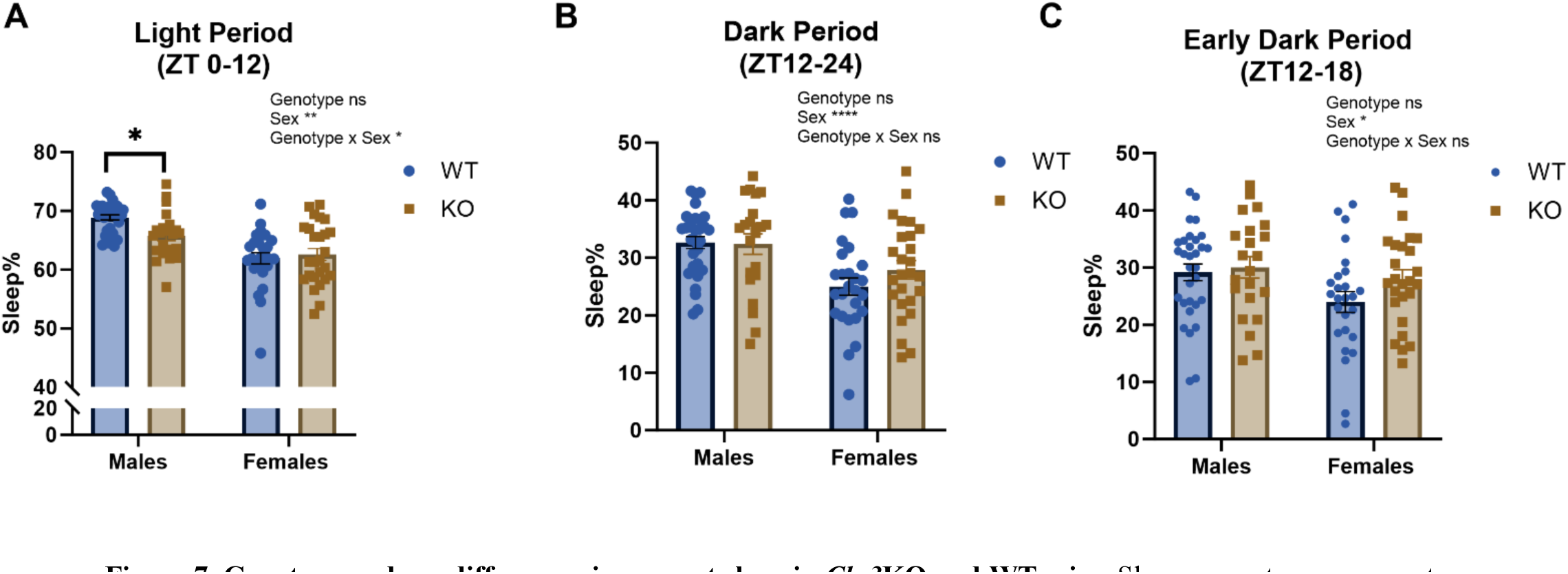
Genotype and sex differences in percent sleep in *Cln3*KO and WT mice. Sleep percent measurements during the LP (ZT0-12), DP (ZT 12-24), and EDP (ZT 12-18) were averaged over days 3-5 of the study were graphed as depicted and analyzed separately. Time over the 24-hour period was shown as ZT, or Zeitgeber Time. Groups included several different ages at the onset of Piezo Sleep acquisition (2 months N= 9 for each genotype; 4 months = 6 months = 9 WT, 7 *Cln3*KO; 8-10 months = 9 WT, 6 *Cln3*KO) and females of several different ages (2 months N=11 for each genotype; 4 months N= 4 WT, 4 *Cln3*KO; 6 months = 6 WT, 5 *Cln3*KO; 8-10 months = 5 WT, 6 *Cln3*KO). (**A**) Percent sleep during the LP in male and female WT and *Cln3*KO groups; (**B**) percent sleep during the DP in male and female WT and *Cln3*KO groups; and (**C**) percent sleep during the EDP in WT and *Cln3*KO male and female mice. Data represented as mean ± SEM (Male WT n= 31; male *Cln3*KO n= 22; female WT n =26; female *Cln3*KO n= 26), *p<0.05, **p<0.001, ****p<0.0001 for overall tests for significance and follow up simple effects tests between groups.

**Table 4.**
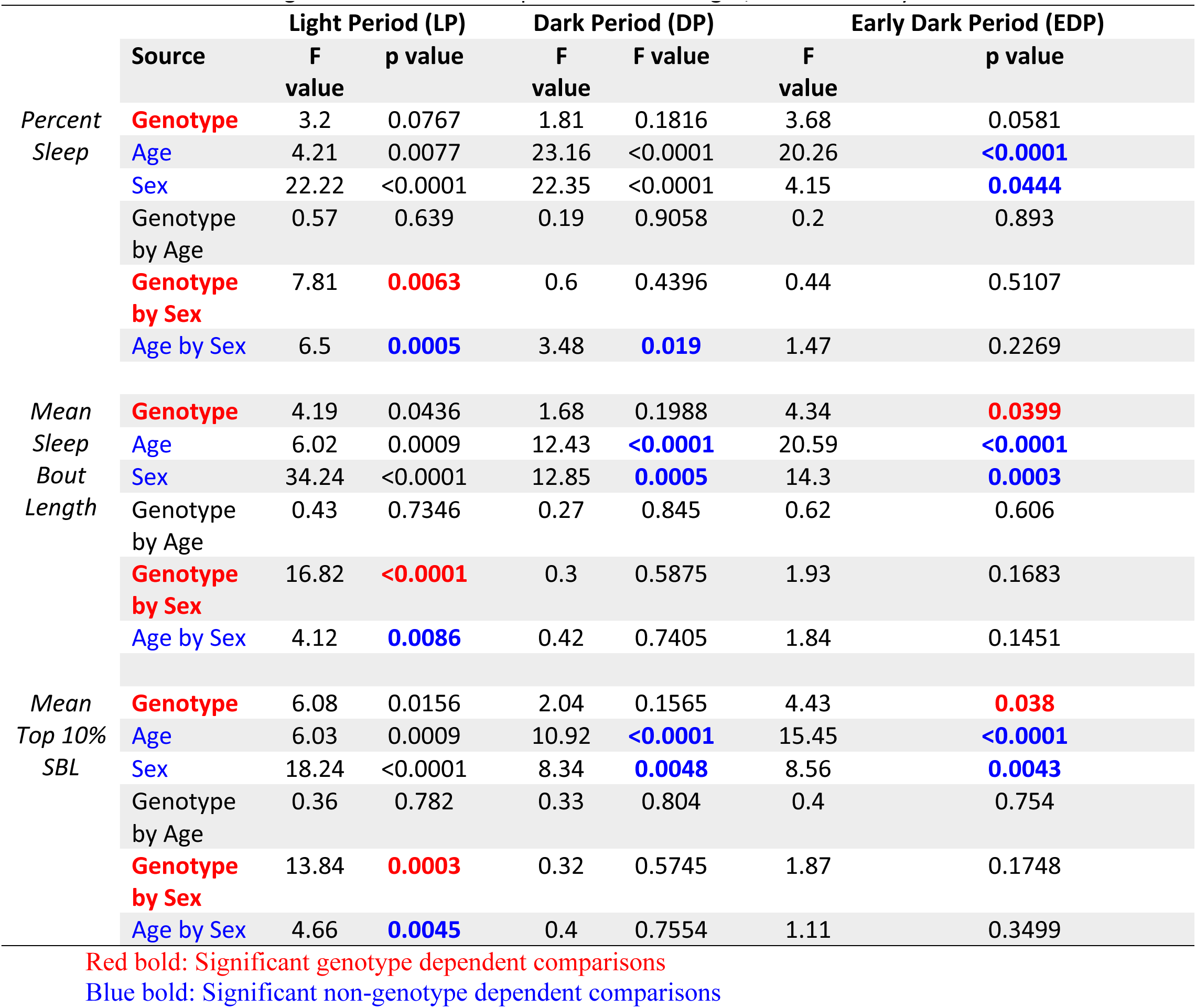
Tests of Significance for the Sleep Differences in Light, Dark and Early Dark Periods

#### Percent sleep sex dimorphism and age dependency

Previous literature has reported sex and age differences in normal sleep of C57BL/6J ^32,33^. This was confirmed in our studies, as there were significant sex effects and age by sex interactions in the PiezoSleep measurements (**Figure S2)**.

Besides genotype by sex interaction in the LP (**Table 4**), percent sleep also displayed the time of day by age by sex interaction over a 24-hour period (**Table 3**), age by sex interactions in both LP and DP (**Table 4**), and sex- and age-dependent differences in EDP (**Table 4**), regardless of genotype.

Specifically, as the time of day by age by sex three-way interaction for percent sleep was significant (p=0.0005; **Table 3**), we followed this up by investigating the time of day by sex interaction within each age group separately. After maintaining the FDR at 0.05, within 2-month-old mice, males had significantly higher levels of sleep percentage during ZT times 1-2, 4, 18, and 19-20.5 (**Figure S2B**), and females had significantly higher levels of sleep percentage during the 13 and 13.5 ZT time points. There were only two significant differences for 4-month-old mice: males had significantly higher levels of sleep percentage during ZT19, and females had a significantly higher percentage during ZT 13 (**Figure S2C**). Six-month males had significantly higher levels of sleep percentage during ZT4.5, 7.5, and 16-20.5, whereas females in the same age range had higher percentages during ZT 13.5-14 (**Figure S2D**). For the oldest mice (8-10 months), males had significantly higher sleep percentages during ZT 0.5-2, 16, 17, 18.5, and 19, whereas females had higher sleep percentages during ZT 13.5 and 23.5 (**Figure S2E**).

As age by sex interactions during both LP and DP were significant (p=0.0005 and 0.019, respectively; **Table 4**), we followed this up and showed that male mice had higher percent sleep than female mice in the 2, 6, and 8-10 months old age groups during the LP (**Figure S3A**) and in the 4- and 6-months old age groups during the DP (**Figure S3B**). In addition, main sex effects during the light (**Figure S3C**), dark (**Figure S3D**), and early dark (**Figure S3E**) periods were all significant (**Table 4**); and main age effects during the light (**Figure S3F**), dark (**Figure S3G**), and early dark (**Figure S3H**) periods were all significant (**Table 4**). Taken together, our results suggest sex dimorphism and age dependency of percent sleep regardless of genotype.

#### Mean sleep bout length abnormalities in Cln3KO mice

For mean sleep bout length, the genotype by time two-way interaction was significant (p=0.0007, **Figure 8A**, **Table 3**). *Cln3*KO mice, as compared to WT, had significantly lower average mean sleep bout lengths during ZT 4 and 7.5 (**Figure 8A**). The genotype by sex two-way interaction was also significant (p=0.0253, **Table 3**). This interaction was driven by male KO mice having a lower average mean sleep bout length than male WT during the LP; female KO mice had higher mean sleep bout lengths during the EDP compared to female WT (**Figure 8B, C**). Therefore, we analyzed mean sleep bout length by dividing the 24 hours into LP and DP/EDP.

**Figure 8.**
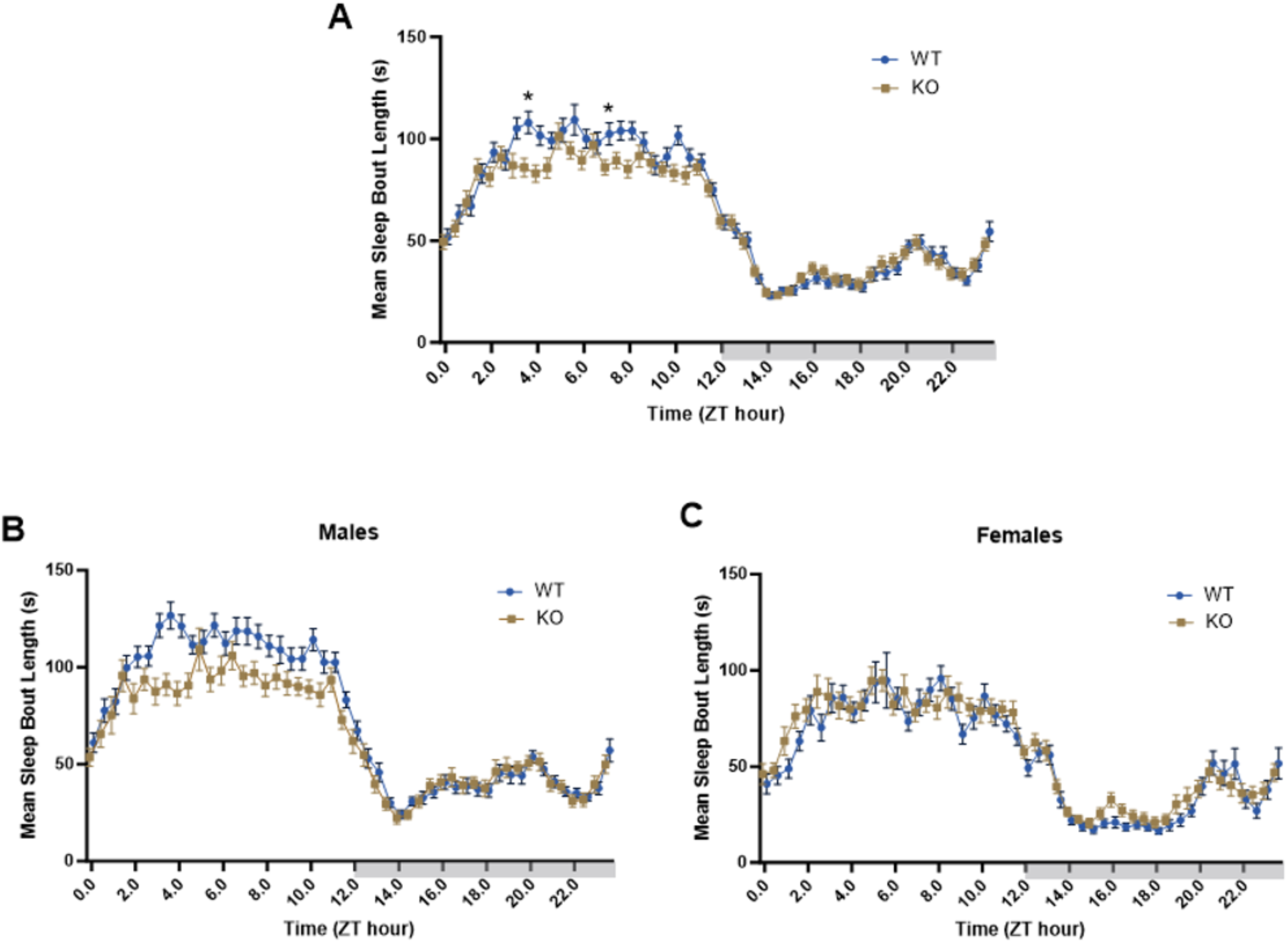
24-hour time course of WT and *Cln3*KO male and female mice. Graphs depict 24-hour sleep traces with mean sleep bout lengths every 30 minutes, averaged from days 3-5. Groups included several different ages at the onset of Piezo Sleep acquisition (2 months N= 9 for each genotype; 4 months = 6 months = 9 WT, 7 *Cln3*KO; 8-10 months = 9 WT, 6 *Cln3*KO) and females of several different ages (2 months N=11 for each genotype; 4 months N= 4 WT, 4 *Cln3*KO; 6 months = 6 WT, 5 *Cln3*KO; 8-10 months = 5 WT, 6 *Cln3*KO). (**A**) WT and *Cln3*KO mice shown in one graph; (**B**) and (**C**) depict WT and *Cln3*KO mice separated by sex. Each data point is represented as mean ± SEM (Male WT n= 31; male *Cln3*KO n= 22; female WT n =26; female *Cln3*KO n= 26), *p<0.05, **p<0.001, ****p<0.0001.

During the LP, for mean sleep bout lengths, genotype by sex interaction was significant (p<0.0001, **Table 4**). A detailed look revealed that during the LP, male *Cln3*KO mice had shorter mean sleep bout lengths compared to WT male mice, while female *Cln3*KO and WT mice showed no differences (**Figure 9A**). During the DP, mean sleep bout length showed insignificant genotype by sex interaction (p=0.5875, **Table 4**) and insignificant main genotype effect (p=0.1988, **Table 4**; **Figure 9B**). However, during the EDP, despite an insignificant genotype by sex interaction (p=0.1683), the main genotype effect reached significance (p=0.0399, **Table 4**). A detailed look revealed that during the EDP, female *Cln3*KO mice seem to have longer mean sleep bout lengths compared to WT female mice, while male WT and *Cln3*KO had similar mean sleep bout lengths (**Figure 9C**).

**Figure 9.**
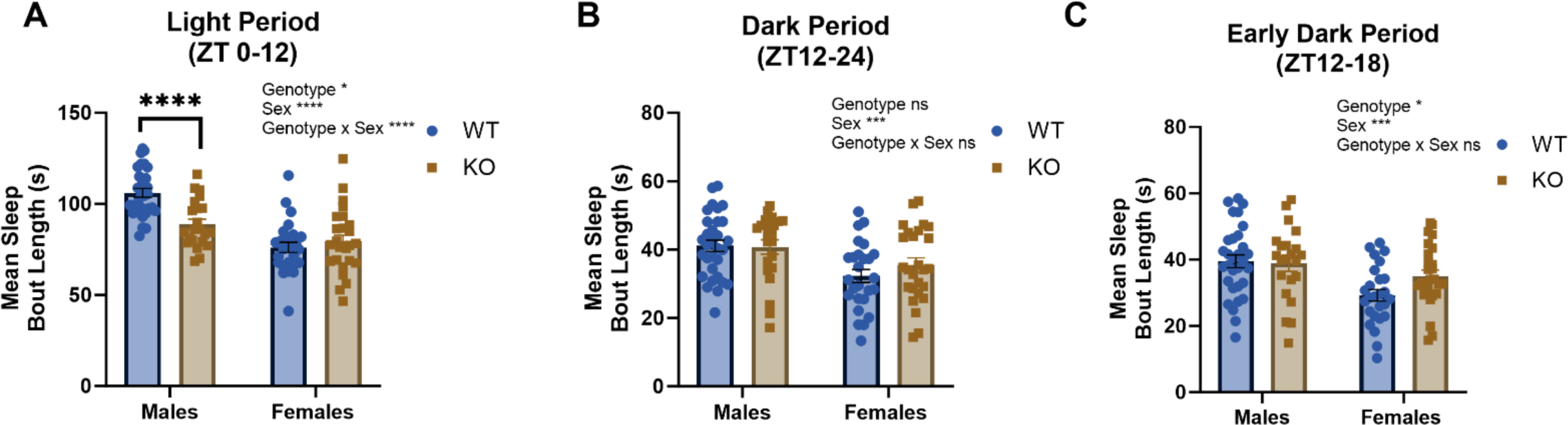
Genotype and Sex Differences in mean sleep bout lengths in WT and *Cln3*KO mice. Bar graphs display mean sleep bout length measurements and were averaged separately as LP (ZT0-12), DP (ZT12-24) and EDP (ZT12-18) over days 3-5 of the study. Groups included several different ages at the onset of Piezo Sleep acquisition (2 months N= 9 for each genotype; 4 months = 6 months = 9 WT, 7 *Cln3*KO; 8-10 months = 9 WT, 6 *Cln3*KO) and females of several different ages (2 months N=11 for each genotype; 4 months N= 4 WT, 4 *Cln3*KO; 6 months = 6 WT, 5 *Cln3*KO; 8-10 months = 5 WT, 6 *Cln3*KO). (**A**) Mean sleep bout lengths during the LP in male and female WT and *Cln3*KO mice; (**B**) DP mean sleep bout lengths for male and female WT and *Cln3*KO mice; and (**C**) Mean sleep bout lengths during the EDP in WT and *Cln3*KO male and female mice. Date represented as mean ± SEM (Male WT n= 31; male *Cln3*KO n= 22; female WT n =26; female *Cln3*KO n= 26), *p<0.05, **p<0.001, ****p<0.0001 for overall tests for significance and follow up simple effects tests between groups.

#### Mean sleep bout length sex dimorphism and age dependency

Besides genotype by sex interaction and genotype by time interaction, mean sleep bout length also displayed the time of day by age by sex interaction over a 24-hour period (**Table 3**), an age by sex interaction in the LP (**Table 4**), and sex- and age-dependent differences in the DP and EDP (**Table 4**), regardless of genotype. Specifically, the time of day by age by sex three-way interaction was significant (p=0.0286, **Table 3**). Thus, the decision was made to follow up this significant interaction by investigating the time of day by sex interaction within each age group separately (**Figure S4B-E**). Maintaining the FDR at 0.05, at 2 months old, male mice had significantly higher average mean sleep bout length than female mice during ZT 1-2, 3, 4, 5.5, 7.5, and 8, 9.5, and 11.5 (**Figure S4B)**. At six months old, males had significantly higher average mean sleep bout lengths than females during ZT 1.5, 2, 4, 4.5, 4, 4.5, 10, and 15; and females had a significantly higher average mean sleep bout length than males during ZT 13.5 (**Figure S4D**). At eight-ten months old, males had significantly higher average mean sleep bout lengths than females during ZT 3, 4.5, 6-7, 8.5, 9, 10.5, and 11 (**Figure S4E**).

Moreover, in the LP, mean sleep bout length also displayed an age by sex interaction (p=0.0086, **Table 4** and **Figure S5A**), and significant sex (p<0.0001, **Table 4** and **Figure S5C**) and age (p=0.0009, **Table 4 and Figure S5F**) effects. In the DP, although age by sex interaction is insignificant, sex effects (p=0.0005, **Table 4** and **Figure S5D**) and age effects (p<0.0001, **Table 4 and Figure S5G**) were significant. In the EDP, sex effects (p=0.0003, **Table 4** and **Figure S5E**) and age effects (p<0.0001, **Table 4** and **Figure S5H**) were significant.

#### Mean Top 10% sleep bout length abnormalities in Cln3KO mice

For mean top 10% sleep bout length, the time of day by genotype by sex three-way interaction was significant (p = 0.0182, **Table 3**). We next analyzed males and females separately, maintaining the FDR at 0.05. Our results show that male *Cln3*KO mice had significantly lower mean top 10% sleep bout lengths during ZT 2.5, 3.5-4.5, 6, 7.5, and 10.5, all of which were during the LP (**Figure 10A**). Although female mice had no significant genotype differences at any time point, female *Cln3*KO mice appeared to have a trend of having higher mean top 10% sleep bout length at ZT time ∼16 (**Figure 10B**). Thus, we next analyzed mean top 10% sleep bout length data in the light, dark, and early dark periods.

**Figure 10.**
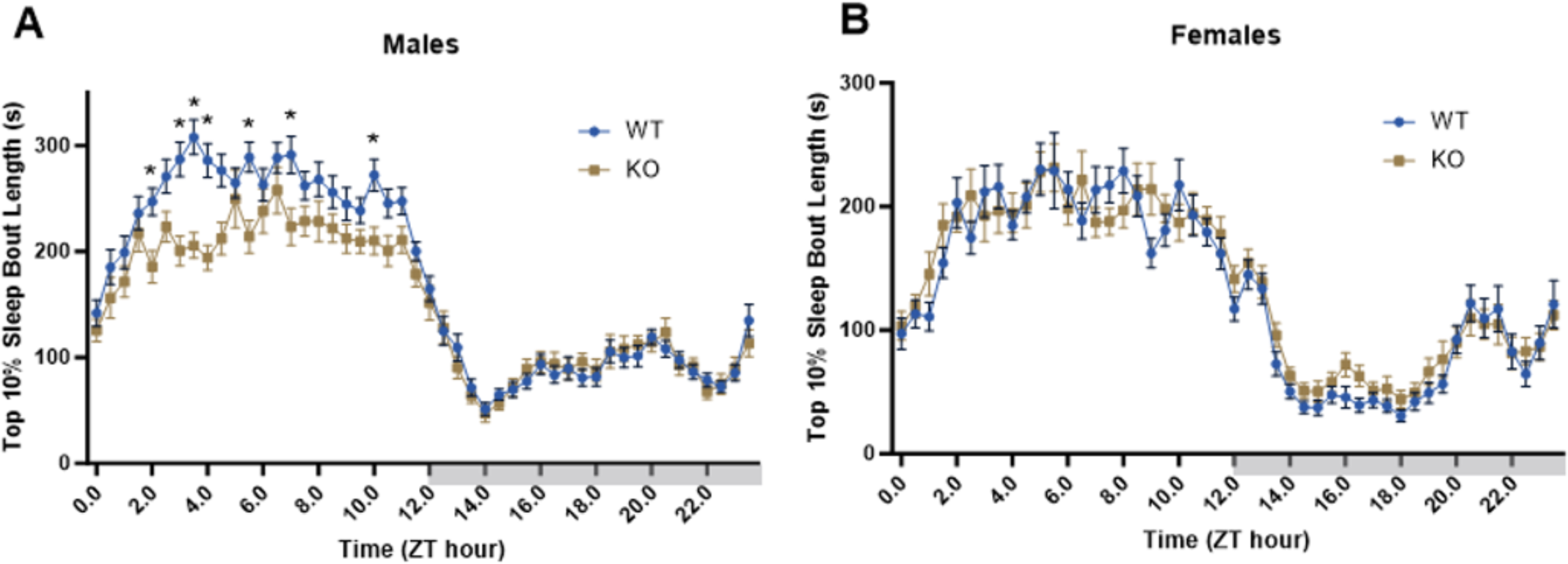
24-hour time course of WT and *Cln3*KO male and female mice. Graphs depict 24-hour sleep traces with top 10% sleep bout lengths every 30 minutes. Groups included several different ages at the onset of Piezo Sleep acquisition (2 months N= 9 for each genotype; 4 months = 6 months = 9 WT, 7 *Cln3*KO; 8-10 months = 9 WT, 6 *Cln3*KO) and females of several different ages (2 months N=11 for each genotype; 4 months N= 4 WT, 4 *Cln3*KO; 6 months = 6 WT, 5 *Cln3*KO; 8-10 months = 5 WT, 6 *Cln3*KO). (**A**) and (**B**) depict WT and Cln3KO mice separated by sex. Each data point is represented as mean ± SEM (Male WT n= 31; male *Cln3*KO n= 22; female WT n =26; female *Cln3*KO n= 26), *p<0.05, **p<0.001, ****p<0.0001.

During the LP, mean top 10% sleep bout length displayed significant genotype by sex interaction (p=0.0003, **Table 4**). A detailed look separating sexes showed that male *Cln3*KO mice had reduced mean top 10% sleep bout lengths compared to WT male, whereas female *Cln3*KO and WT mice showed no significant differences (**Figure 11A**). During the DP, mean top 10% sleep bout length displayed insignificant genotype by sex interaction (p=0.5745, **Table 4**), whereas main genotype effect was approaching significance (p=0.1565, **Table 4**). All four groups were similar in this sleep measurement during this 12-hour phase. However, during the EDP, genotype by sex interaction was approaching significance (p=0.1748, **Table 4**) and contrary to other findings, regardless of sex, *Cln3*KO mice had significantly higher mean top 10% sleep bout lengths than WT (p=0.038, **Table 4** and **Figure 11D**). A detailed look separating sexes showed that during the EDP, female *Cln3*KO mice had increased mean top 10% sleep bout lengths compared to female WT mice whereas male *Cln3*KO and WT were similar (**Figure 11C**).

**Figure 11.**
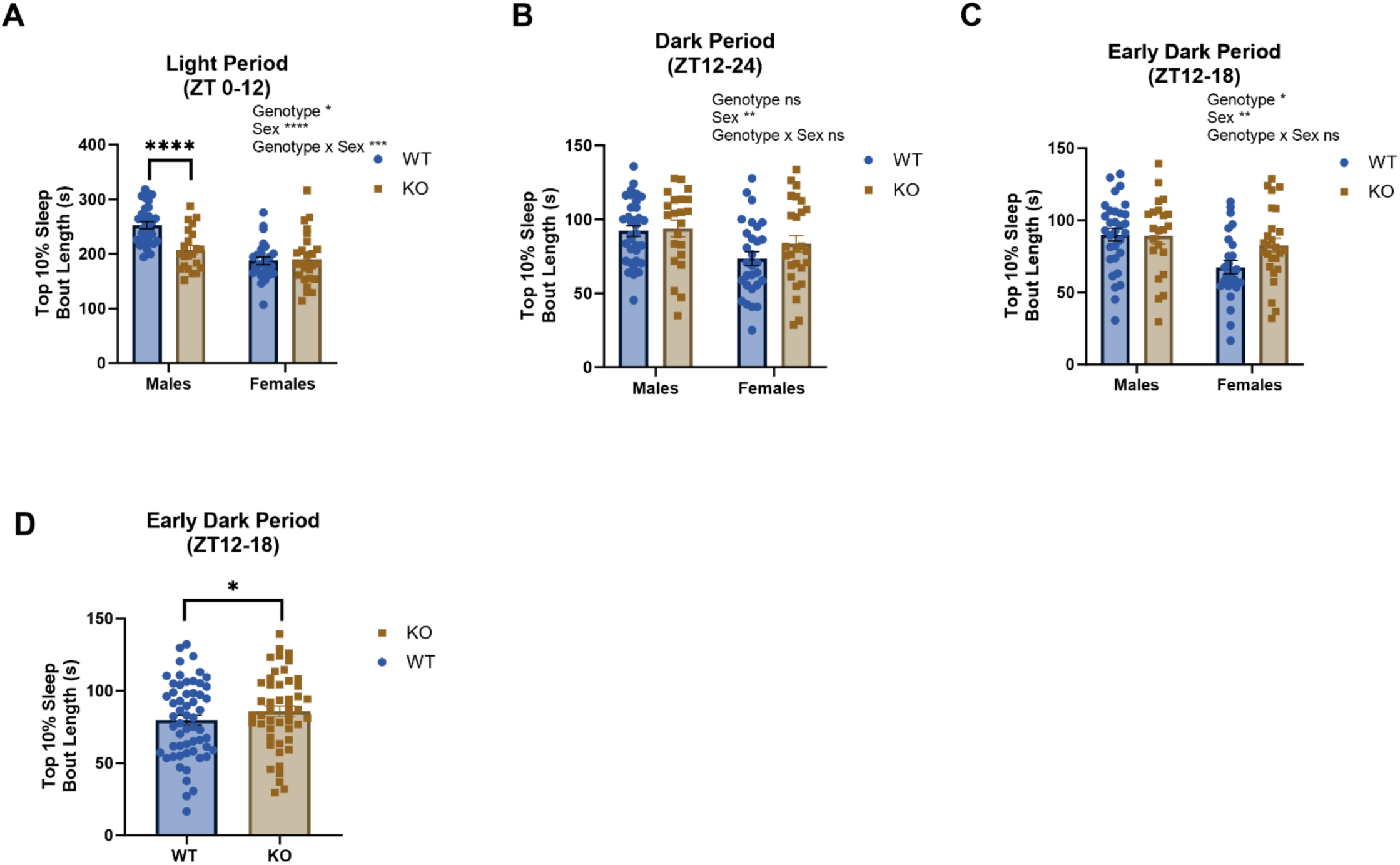
Genotype and Sex Differences in top 10% sleep bout lengths in WT and *Cln3*KO mice. Bar graphs display mean sleep bout length measurements and were averaged separately as LP (ZT0-12), DP (ZT12-24) and EDP (ZT12-18) over days 3-5 of the study. Groups included several different ages at the onset of Piezo Sleep acquisition (2 months N= 9 for each genotype; 4 months = 6 months = 9 WT, 7 *Cln3*KO; 8-10 months = 9 WT, 6 *Cln3*KO) and females of several different ages (2 months N=11 for each genotype; 4 months N= 4 WT, 4 *Cln3*KO; 6 months = 6 WT, 5 *Cln3*KO; 8-10 months = 5 WT, 6 *Cln3*KO). (**A**) Top 10% sleep bout lengths during the LP in male and female WT and *Cln3*KO mice; (**B**) DP top 10% sleep bout lengths for male and female WT and *Cln3*KO mice; and (**C**) EDP top 10% sleep bout lengths for male and female WT and *Cln3*KO mice; (**D**) EDP top 10% sleep bout lengths for WT and *Cln3*KO mice. Date represented as mean ± SEM (Male WT n= 31; male *Cln3*KO n= 22; female WT n =26; female *Cln3*KO n= 26), *p<0.05, **p<0.001, ****p<0.0001 for overall tests for significance and follow up simple effects tests between groups.

#### Mean top 10% sleep bout length sex dimorphism and age dependency

Besides the time of day by genotype by sex three-way interaction, mean top 10% sleep bout lengths also displayed the time of day by age by sex interaction over a 24-hour period (**Table 3**), an age by sex interaction in the LP (p=0.0003, **Table4**), and sex- and age-dependent differences in both the dark and early dark periods (**Table 4**), regardless of genotype.

Specifically, the time of day by age by sex three-way interaction was significant (p < 0.0001, **Table 3**). To follow up this significant interaction, we investigated the time of day by sex interaction within each age group separately. After maintaining the FDR at 0.05, 2-month-old male mice had significantly higher mean top 10% sleep bout lengths during ZT 1-2, 3, 4, 11.5, and 20 (**Figure S6B**). Six-month males had significantly higher mean top 10% sleep bout lengths during ZT 4-5, 7, 7.5, and 19, but significantly lower mean top 10% sleep bout lengths during ZT 13.5 (**Figure S6D**). For the 8-10 months old mice, males had significantly higher mean top 10% sleep bout lengths during ZT 4.5, 6-7, 8.5, 9, 10.5, 11, and 12.5 (**Figure S6E**).

Moreover, in the LP, mean top 10% sleep bout length also displayed an age by sex interaction (p=0.0045, **Table 4** and **Figure S6A**) as well as significant age effects (p=0.0009, **Table 4** and **Figure S7F**). The DP displayed significant sex effects (p=0.0048, **Table 4** and **Figure S7B, D**) and age effects (p<0.0001, **Table 4** and **Figure S7G**). The EDP also showed significant sex effects (p=0.0043, **Table 4** and **Figure S7E**) and age effects (p<0.0001, **Table 4** and **Figure S7H**).

## Discussion

### Validity of the PiezoSleep system for monitoring sleep abnormalities in a mouse model of CLN3 disease

We chose to use the noninvasive PiezoSleep system to monitor sleep, as the gold standard EEG/EMG method is labor intensive and requires extensive instrumentation, thus making it difficult to assess a large cohort of animals. To ensure the accuracy and reliability of our findings using PiezoSleep, we validated it against the gold standard EEG/EMG method. Our results demonstrate high Spearman rank correlation coefficients (70% or above; **Figure 2A**), high AUC values (>90%) in ROC analysis (**Figure 2C**), and strong and consistent correlations between the PiezoSleep decision statistic and mean-squared EMG power (**Figure S1**), indicating strong agreement between PiezoSleep and the EEG/EMG method for discriminating between sleep and wake states. These results confirm the reliability of PiezoSleep in assessing sleep architecture across different genotypes and periods of the LD cycle, with sensitivity and specificity exceeding 90%.

While sleep% estimates were highly consistent between the two methods, discrepancies were observed in sleep MBL, with EMG resulting in longer MBL compared to PiezoSleep (**Figure 3**). This difference may be attributed to the piezo sensor’s sensitivity to brief arousals in sleep, particularly during REM sleep, potentially due to its location on the animal’s ventral surface. This difference does not affect the results when comparing different genotype, sex, or age, due to high Spearman rank correlation coefficients (70% or above; **Figure 2A**). Our analysis further demonstrated that after discounting brief arousals, the piezo and EMG estimates of MBL converged (**Figure S1**), indicating overall consistency between the two methods.

### Comparison of sleep data in CLN3 disease mouse models versus human case studies

Clinical data has reported sleep disturbances as the most common symptoms experienced by juvenile ceroid lipofuscinosis patients ^17–19,21,34^. However, possible sleep abnormalities have not previously been investigated in animal models of CLN3 disease. It was reported that starting at 6 months of age, *Cln3^Δex^*^7–8^*^/Δex^*^7–8^, the CLN3 disease mouse model with a ∼1-kb deletion that precisely mimics the most common human mutation, showed increased spiking activity on EEG recordings in both the cortex and hippocampus ^35^. Quantification of power in the EEG frequency bands also revealed a decrease in low frequency activity in *Cln3^Δex^*^7–8^*^/Δex^*^7–8^ compared to WT mice ^35^. While this immediately suggests a reduction in slow wave sleep in the *Cln3^Δex^*^7–8^*^/Δex^*^7–8^ mice, no sleep analysis was done in that study to investigate the possibility and the correlation of sleep differences, if any, with spiking activity.

Our current study aiming at filling in the knowledge gap and characterizing sleep-wake alterations in *Cln3^Δex^*^1–6^*^/Δex^*^1–6^ (*Cln3*KO) at ages when pathologies associated with CLN3 disease (e.g., intracellular accumulation of auto fluorescent material) are present ^22^ is, to our knowledge, the first study on sleep disturbances of CLN3 disease animal models. Our PiezoSleep data show that sleep percentages, mean sleep bout lengths, and top 10% sleep bout lengths are reduced in male *Cln3*KO (compared to WT) mice during the LP. As mice are nocturnal animals, and therefore less active in the LP, our findings of sleep disturbances in the male *Cln3*KO mice recapitulate disrupted sleep during the night in CLN3 disease patients ^17,21^. The female *Cln3*KO mice showed an increase in sleep percentage, mean sleep bout length, and top 10% sleep bout length in the EDP. As the DP is the active period for mice, our findings align with daytime sleepiness documented in CLN3 disease patients ^17^.

In this current study, we observed sex differences in the characteristics of sleep abnormalities of *Cln3*KO mice: female *Cln3*KO mice seemed to show subtle differences from their WT counterparts during the EDP, whereas male *Cln3*KO mice were more severely affected during the LP. In contrast, literatures on sex differences in clinical patients report mixed information. It was suggested that females may develop symptoms on average slightly later and die one year later than males, but experience more severe symptoms compared to males ^36,37^; nevertheless, assessment of sleep was lacking in these studies. Future investigations of sex differences in the sleep disturbances of both CLN3 disease patients and mouse models are needed to better understand the sleep disturbances associated with the disease and mechanisms behind these symptoms.

### Caveats and future directions

Although we have shown sex dimorphism in the characteristics of sleep abnormalities of *Cln3*KO (*Cln3^Δex^*^1–6^*^/Δex^*^1–6^) mice across different ages, we have not tested if other CLN3 disease mouse models, such as *Cln3^Δex^*^7–8^*^/Δex^*^7–8^, also exhibit similar sex-dependent sleep abnormalities.

Although a small sample analysis with EEG/EMG validated the accuracy of the PiezoSleep system in monitoring sleep abnormalities in CLN3 disease and likely other neurodegenerative disease mouse models, differences in EEG/EMG-based sleep metrics between *Cln3*KO and WT female mice did not reach statistical significance due to the small sample size (**Figures 4 & 5**), except for a trend in female *Cln3*KO towards longer MBL than female WT for REM sleep in DP (**Figure 5F**). An EEG/EMG study with a larger cohort is needed to obtain more conclusive information regarding disturbances in sleep composition and architecture in *Cln3*KO.

Moreover, the mechanisms causing sleep abnormalities in the *Cln3*KO mouse model warrants further investigation. The role CLN3 plays is yet to be fully elucidated, but the pathologies observed, particularly in neurons, that arise from a loss of CLN3 could also be explored in relation to sleep abnormalities. Accumulation of ATP mitochondrial subunit c, along with other proteins and lipids, have been reported and are considered pathological hallmarks of the disease ^38^. Neuronal cell death has been observed in CLN3 disease patients, resulting in reduced hippocampal and cortical volume ^39,40^. Moreover, consistent with the previous report ^35^, we also found increased spiking activity in *Cln3*KO (data not shown). Modulation of spiking activity by sleep stage will suggest potential contribution of spiking to impaired sleep quality in *Cln3*KO but would require an EEG/EMG analysis in a larger sample. Also. whether the sleep disturbances reported here in the *Cln3*KO mouse model are due to circadian rhythm abnormalities is yet to be investigated, even though in human CLN3 disease patients, sleep fragmentation and sleep phase irregularities could not be explained by abnormalities in the circadian regulatory system ^19^.

Conversely, whether vision loss and the cognitive and behavioral impairments that patients suffer from are associated with and may be further exacerbated by sleep disturbances is unknown. Poor sleep may result in cognition, learning, and memory disfunctions, and has been linked with other neurogenerative diseases, such as Alzheimer’s disease ^41^. Are the sleep disturbances in CLN3 disease contributing to progression of the disease? Indeed, chronic sleep disruptions can pose a serious impact on brain health, and in patients suffering from CLN3 disease, low-quality sleep could be a potential driver of neurologic impairments and decline associated with the disease.

The sleep phenotype obtained from this study will be useful for future therapeutic development. Exploring treatment strategies to potentially recuperate sleep in CLN3 disease patients could slow the progression of disease symptoms and improve the quality of life for these individuals.

## Acknowledgement

**Support:** Q.J.W. thanks Marilyn Duncan for allowing the usage of PiezoSleep equipment. This project was supported by the University of Kentucky Neuroscience Research Priority Area (NRPA) pilot grants and the NIH National Center for Research Resources and the National Center for Advancing Translational Sciences grant UL1TR001998 (Center for Clinical and Translational Science High Impact pilot grant) to QJW; and NIH R42NS107148 to SS.

## Supplementary Figures

**Figure S1.**
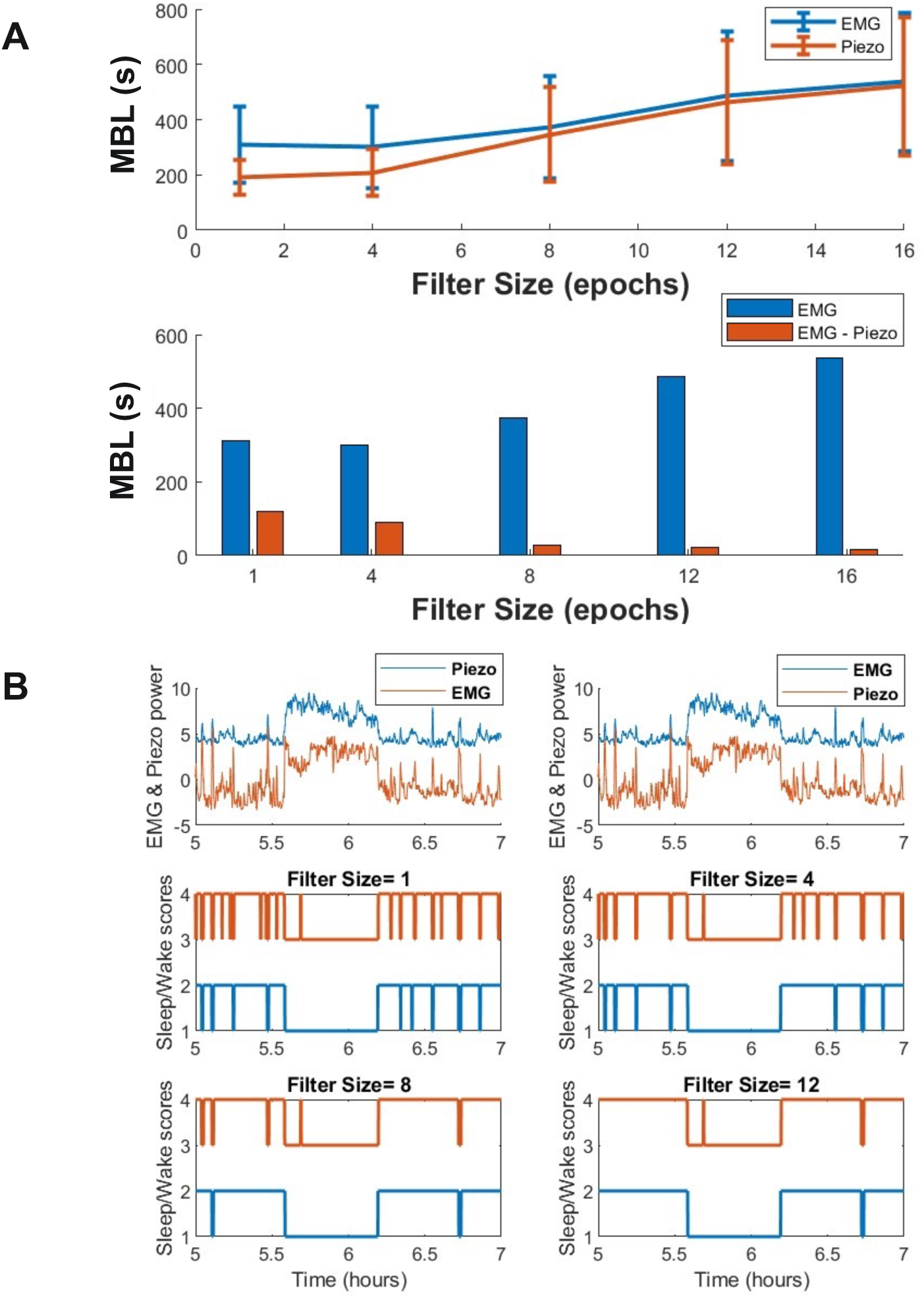
Convergence of EMG and piezo sleep metrics upon progressive removal of brief arousals. (**A**) Demonstration of the convergence of EMG and piezo sleep MBL and a trend toward zero difference as brief arousals are progressively removed by merging adjacent sleep bouts; (**B**) A sample plot that shows the corresponding sleep scores at various levels of brief arousal removal, providing visual insight into the convergence of EMG and piezo sleep metrics.

**Figure S2.**
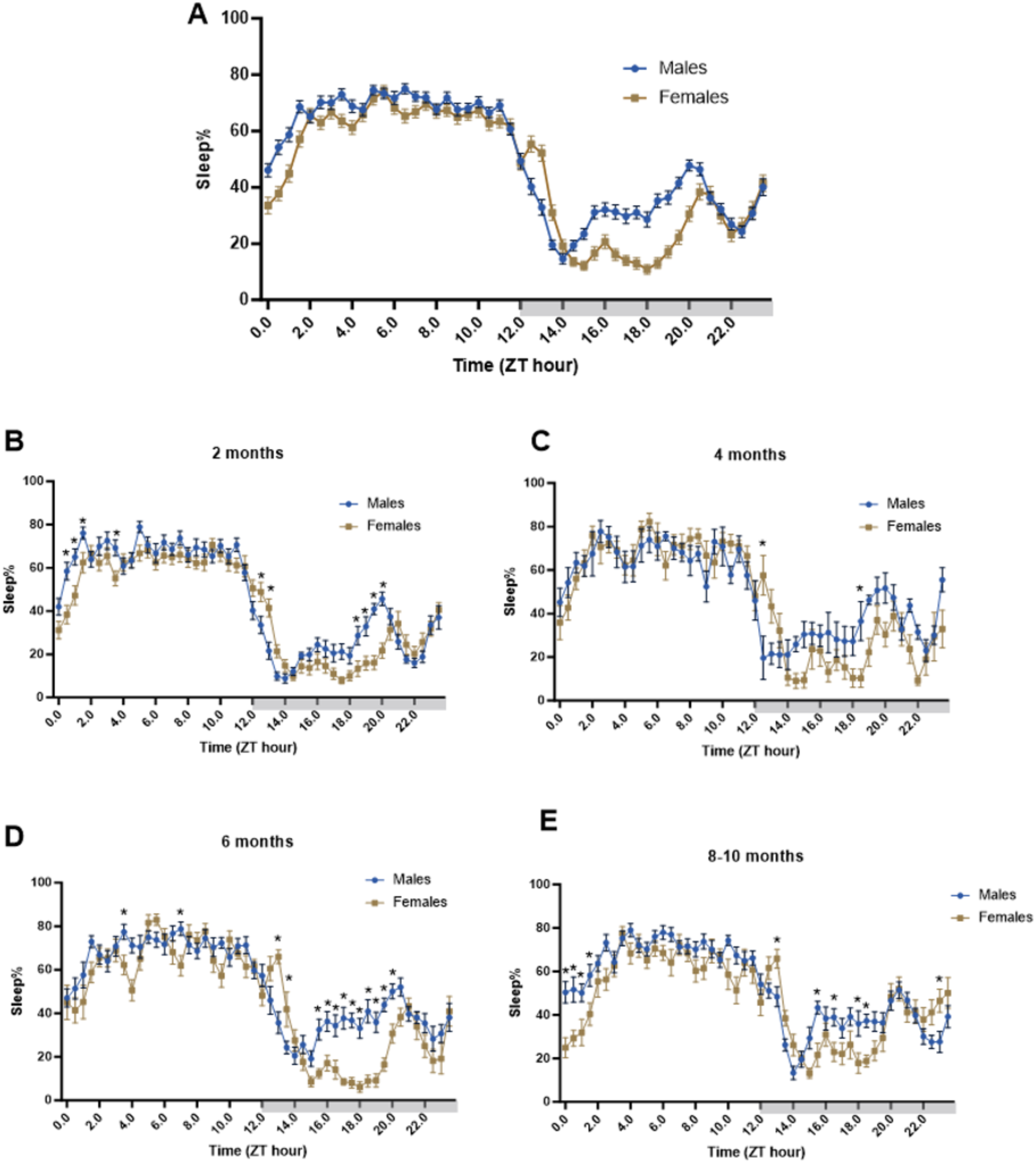
Age by Sex differences in sleep percentage 24-hour time courses. Graphs depict 24-hour sleep traces with sleep percentages every 30 minutes, averaged from days 3-5. Groups included several different ages at the onset of Piezo Sleep acquisition (2 months N= 9 for each genotype; 4 months = 6 months = 9 WT, 7 *Cln3*KO; 8-10 months = 9 WT, 6 *Cln3*KO) and females of several different ages (2 months N=11 for each genotype; 4 months N= 4 WT, 4 *Cln3*KO; 6 months = 6 WT, 5 *Cln3*KO; 8-10 months = 5 WT, 6 *Cln3*KO). (**A**) Male and females represented in one graph; (**B-E**) show 2-month, 4-month, 6-month, and 8-10-month age groups separated by sex. Each data point is represented as mean ± SEM, *p<0.05, **p<0.001, ****p<0.0001.

**Figure S3.**
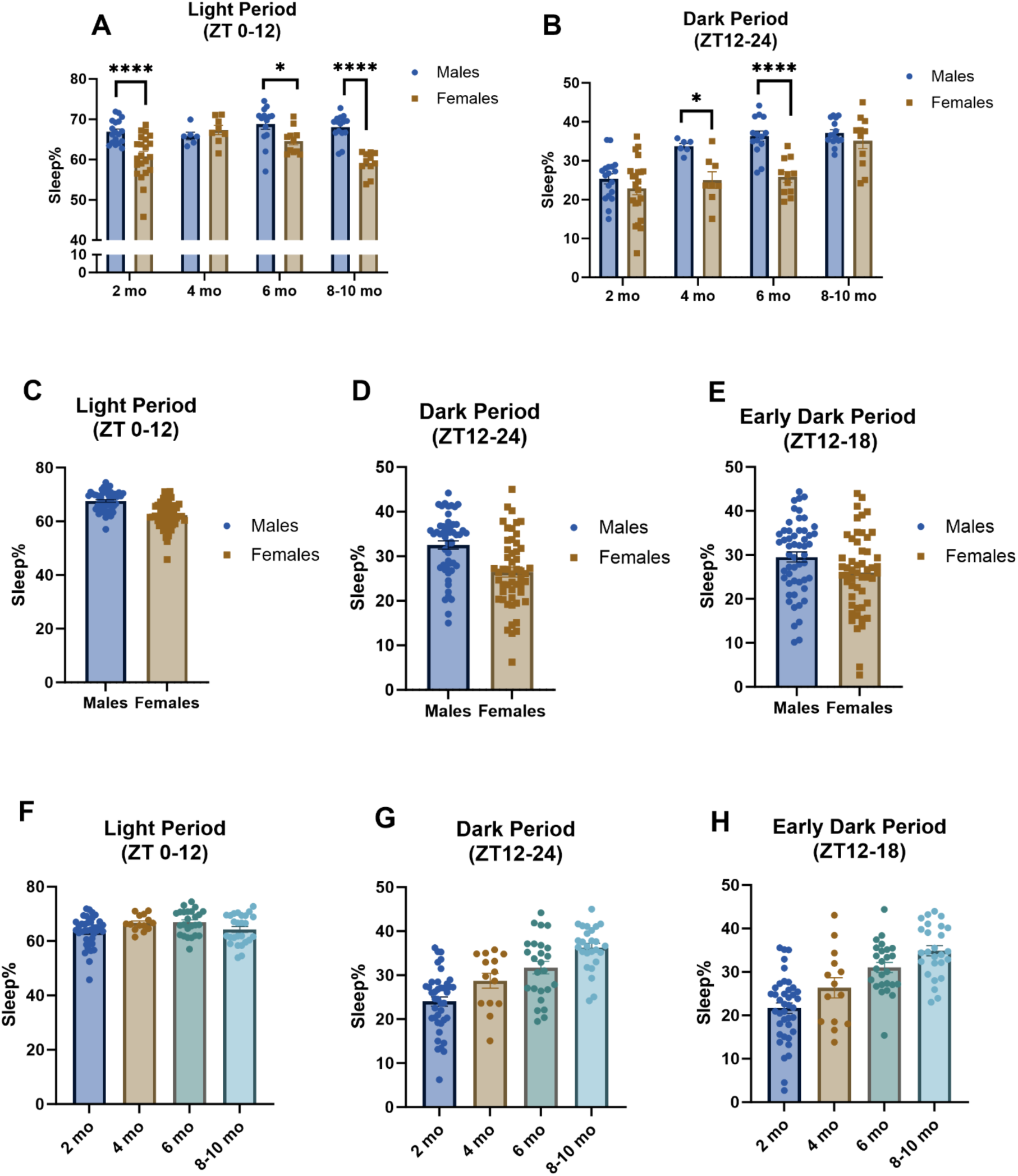
Age by sex differences in sleep percentages in light and dark periods. Bar graphs represent LP (ZT 0-12), DP (ZT 12-24), and EDP (ZT 12-18) sleep percentages averaged over days 3-5 of the study. Groups included several different ages at the onset of Piezo Sleep acquisition (2 months N= 9 for each genotype; 4 months = 6 months = 9 WT, 7 *Cln3*KO; 8-10 months = 9 WT, 6 *Cln3*KO) and females of several different ages (2 months N=11 for each genotype; 4 months N= 4 WT, 4 *Cln3*KO; 6 months = 6 WT, 5 *Cln3*KO; 8-10 months = 5 WT, 6 *Cln3*KO). (**A**) LP sleep percent with male and female separated into their respective age groups (2, 4, 6, 8-10 months); (**B**) DP sleep percentages with male and female separated by age; (**C-E**) LP, DP, and EDP graphs with mice only separated by sex; (**F-H**) LP, DP, and EDP graphs with mice only separated by age. Data represented as mean ± SEM, *p<0.05, **p<0.001, ****p<0.0001 for follow-up simple effects tests.

**Figure S4.**
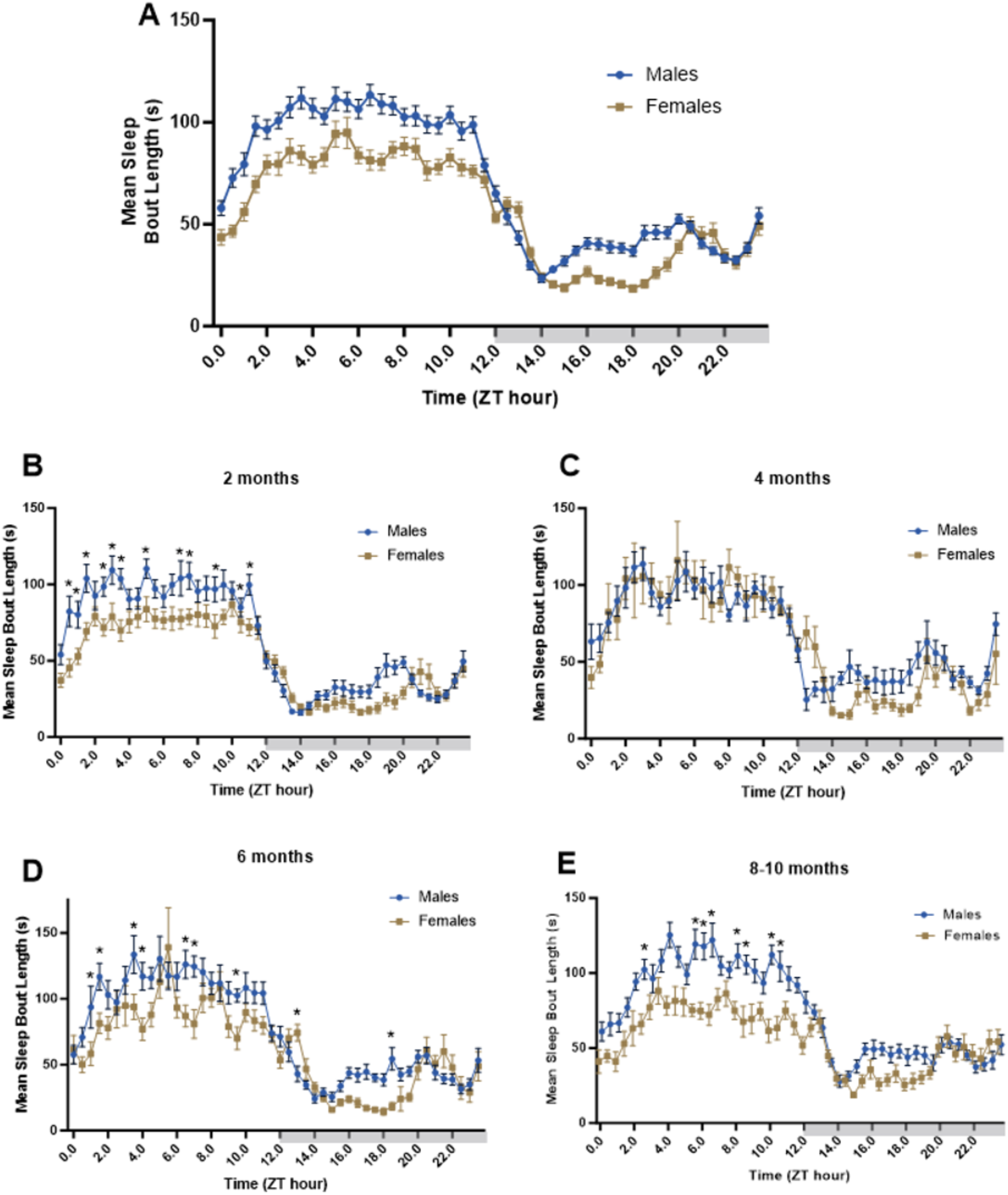
Age by Sex differences in mean sleep bout length 24-hour time courses. Graphs depict 24-hour sleep traces with mean sleep bout lengths every 30 minutes, averaged from days 3-5. Groups included several different ages at the onset of Piezo Sleep acquisition (2 months N= 9 for each genotype; 4 months = 6 months = 9 WT, 7 *Cln3*KO; 8-10 months = 9 WT, 6 *Cln3*KO) and females of several different ages (2 months N=11 for each genotype; 4 months N= 4 WT, 4 *Cln3*KO; 6 months = 6 WT, 5 *Cln3*KO; 8-10 months = 5 WT, 6 *Cln3*KO). (**A**) Male and females represented in one graph. (**B-E**) show 2-month, 4-month, 6-month, and 8-10-month age groups separated by sex. Each data point is represented as mean ± SEM.

**Figure S5.**
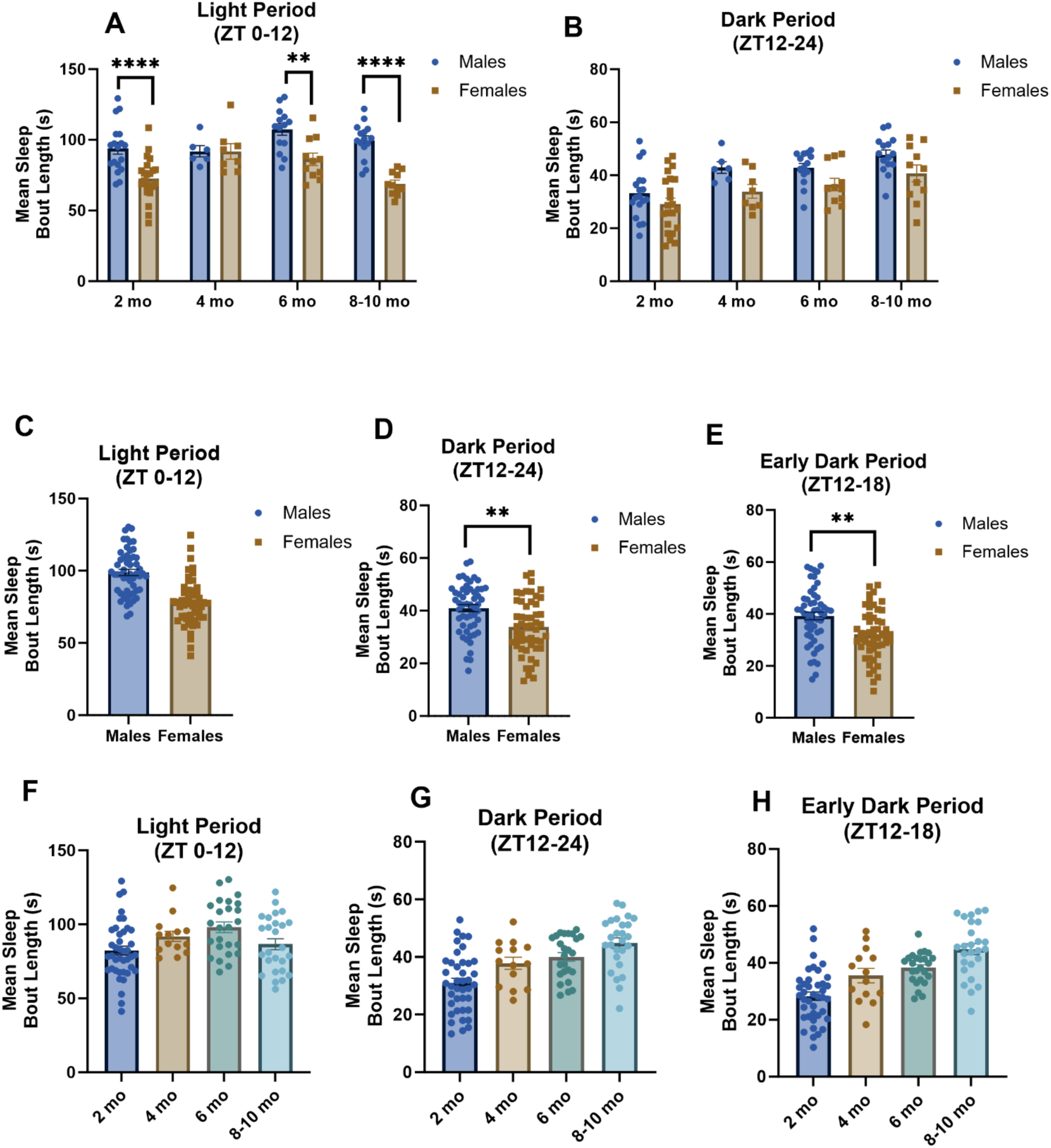
Age by sex differences in mean sleep bout lengths in light and dark periods. Bar graphs represent LP (ZT 0-12), DP (ZT 12-24), and EDP (ZT 12-18) mean sleep bout lengths averaged over days 3-5 of the study. Groups included several different ages at the onset of Piezo Sleep acquisition (2 months N= 9 for each genotype; 4 months = 6 months = 9 WT, 7 *Cln3*KO; 8-10 months = 9 WT, 6 *Cln3*KO) and females of several different ages (2 months N=11 for each genotype; 4 months N= 4 WT, 4 *Cln3*KO; 6 months = 6 WT, 5 *Cln3*KO; 8-10 months = 5 WT, 6 *Cln3*KO). (**A**) LP sleep percent with male and female separated into their respective age groups (2, 4, 6, 8-10 months); (**B**) DP mean sleep bout lengths with male and female separated by age; (**C-E**) LP, DP, and EDP bar graphs with mice only separated by sex; (**F-H**) LP, DP, and EDP graphs with mice only separated by age. Data represented as mean ± SEM, *p<0.05, **p<0.001, ****p<0.0001.

**Figure S6.**
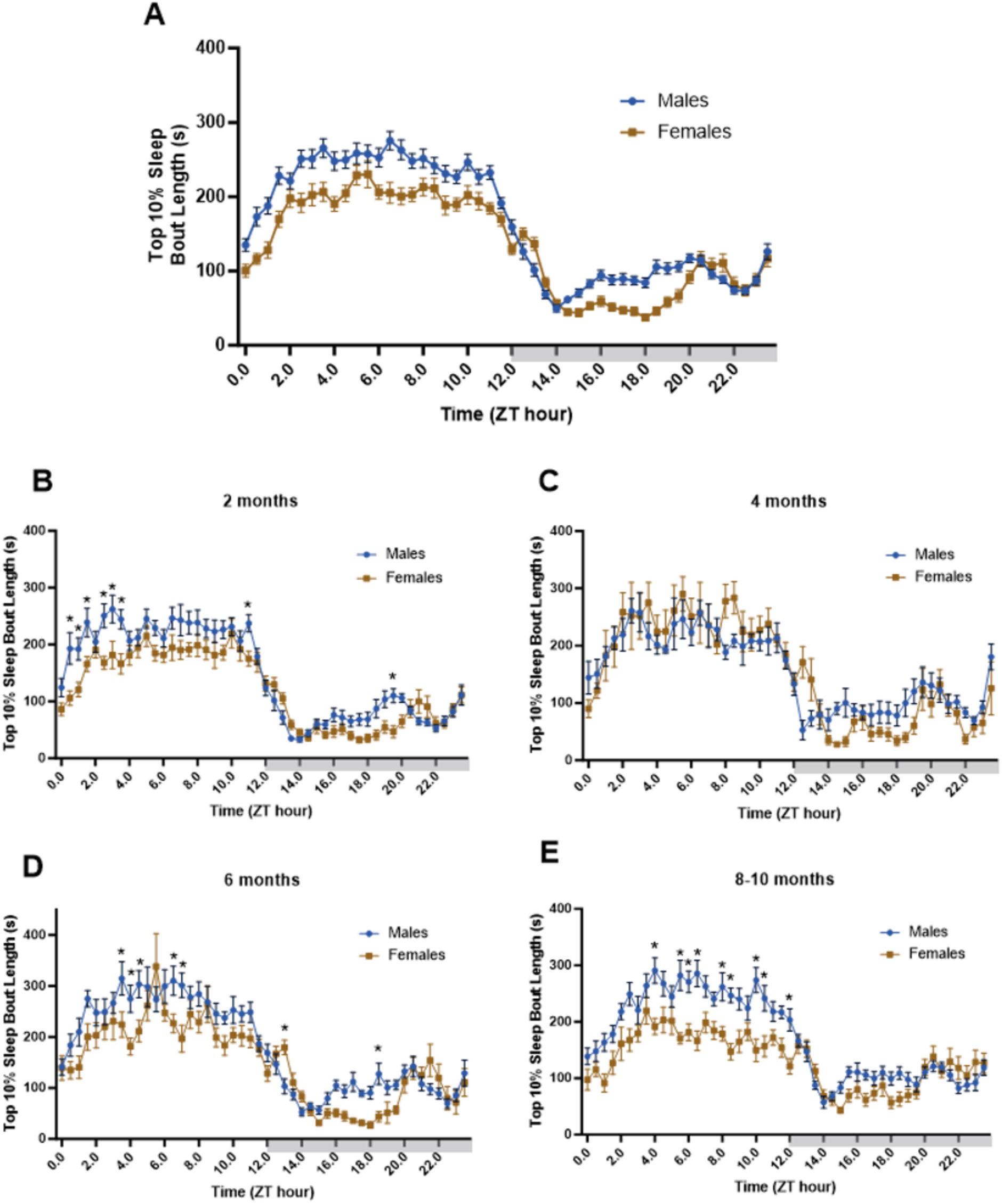
Age by Sex differences in top 10% sleep bout length 24-hour time courses. Graphs depict 24-hour sleep traces with top 10% sleep bout lengths every 30 minutes, averaged from days 3-5. (**A**) Male and females represented in one graph. Groups included several different ages at the onset of Piezo Sleep acquisition (2 months N= 9 for each genotype; 4 months = 6 months = 9 WT, 7 *Cln3*KO; 8-10 months = 9 WT, 6 *Cln3*KO) and females of several different ages (2 months N=11 for each genotype; 4 months N= 4 WT, 4 *Cln3*KO; 6 months = 6 WT, 5 *Cln3*KO; 8-10 months = 5 WT, 6 *Cln3*KO). (**B-E**) show 2-month, 4-month, 6-month, and 8-10-month age groups separated by sex. Each data point is represented as mean ± SEM.

**Figure S7.**
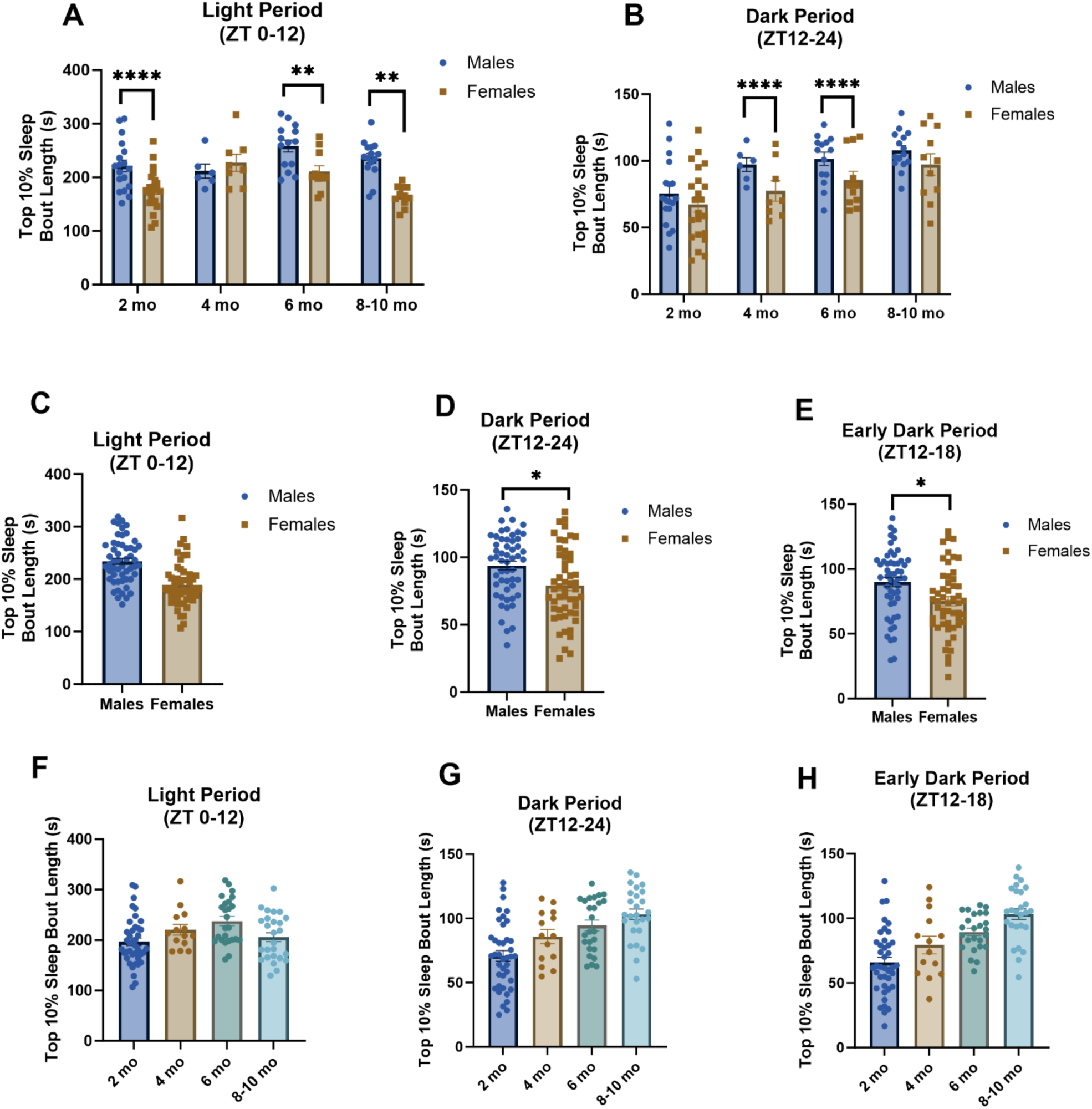
Age by sex differences in top 10% sleep bout lengths in light and dark periods. Bar graphs represent LP (ZT 0-12), DP (ZT 12-24), and EDP (ZT 12-18) top 10% sleep bout lengths averaged over days 3-5 of the study. Groups included several different ages at the onset of Piezo Sleep acquisition (2 months N= 9 for each genotype; 4 months = 6 months = 9 WT, 7 *Cln3*KO; 8-10 months = 9 WT, 6 *Cln3*KO) and females of several different ages (2 months N=11 for each genotype; 4 months N= 4 WT, 4 *Cln3*KO; 6 months = 6 WT, 5 *Cln3*KO; 8-10 months = 5 WT, 6 *Cln3*KO). (**A**) LP top 10% sleep bout lengths with male and female separated into their respective age groups (2, 4, 6, 8-10 months); (**B**) DP top 10% sleep bout lengths with male and female separated by age; (**C-E**) LP, DP, and EDP bar graphs with mice only separated by sex; (**F-H**) LP, DP, and EDP graphs with mice only separated by age. Data represented as mean ± SEM, *p<0.05, **p<0.001, ****p<0.0001.

